# A spatially resolved single cell atlas of human gastrulation

**DOI:** 10.1101/2020.07.21.213512

**Authors:** Richard C.V. Tyser, Elmir Mahammadov, Shota Nakanoh, Ludovic Vallier, Antonio Scialdone, Shankar Srinivas

**Affiliations:** Department of Physiology, Anatomy and Genetics, South Parks Road, University of Oxford, Oxford, OX1 3QX, UK; Institute of Epigenetics and Stem Cells, Helmholtz Zentrum München – German Research Center for Environmental Health, Munich, 81377, Germany; Institute of Functional Epigenetics, Helmholtz Zentrum München – German Research Center for Environmental Health, Neuherberg, 85764, Germany; Institute of Computational Biology, Helmholtz Zentrum München – German Research Center for Environmental Health, Neuherberg, 85764, Germany; Wellcome - MRC Cambridge Stem Cell Institute, Jeffrey Cheah Biomedical Centre, Puddicombe Way, Cambridge Biomedical Campus, Cambridge, CB2 0AW, UK

## Abstract

Gastrulation is the fundamental process during the embryogenesis of all multicellular animals through which the basic body plan is first laid down. It is pivotal in generating cellular diversity coordinated with spatial patterning. Gastrulation in humans occurs in the third week following fertilization. Our understanding of this process in humans is extremely limited, and based almost entirely on experimental models. Here, we characterize in a spatially resolved manner the single cell transcriptional profile of an entire gastrulating human embryo approximately 16 to 19 days after fertilization. We used these data to provide the first unequivocal demonstration that human embryonic stem cells represent the early post implantation epiblast. We identified both primordial germ cells and red blood cells, which had never been characterized so early during human development. Comparison with mouse gastrula transcriptomes revealed many commonalities between the human and mouse but also several key differences, particularly in FGF signaling, that we validated experimentally. This unique dataset offers a unique glimpse into a central but generally inaccessible stage of our development, provides new context for interpreting experiments in other model systems and represents a valuable resource for guiding directed differentiation of human cells *in vitro*.

## INTRODUCTION

Gastrulation is a fundamental process during embryonic development, conserved across all multicellular animals. It is characterized by large scale morphogenetic remodelling that leads to the conversion of an early pluripotent embryonic cell layer into the three primary ‘germ layers’ typical of the majority of metazoans: an outer ectoderm, inner endoderm and intervening mesoderm layer. The morphogenesis of these three layers of cells is closely coordinated with cellular diversification, laying the foundation for the generation of the hundreds of distinct specialized cell types in the animal body. The process of gastrulation has attracted tremendous attention in a broad range of experimental systems ranging from worms, flies, echinoderms, fish, chick, rabbits and mice to name just a few (Stern 2004; Briggs et al. 2018; Wagner et al. 2018; Nowotschin et al. 2019; Pijuan-Sala et al. 2019). However our understanding of gastrulation in humans is based almost entirely on extrapolation from these model systems and from limited collections of fixed whole samples and histological sections of human gastrulae (O’Rahilly and Müller 2010; Yamaguchi and Yamada 2019; Florian and Hill 1935; De Bakker et al. 2016), some of which date back to over a century ago. In humans the process of gastrulation starts approximately 14 days after fertilization and continues for slightly over a week. Donations of human fetal material at these early stages, when many people might not even know they are pregnant, are exceptionally rare, making it nearly impossible to study gastrulation directly. Therefore, efforts to understand human gastrulation have predominantly focused on *in vitro* models such as monolayers of human Embryonic Stem Cells (hESCs) cultured on circular micropatterns (Warmflash et al. 2014). More recently, these have been extended to hESC colonies engrafted into chick embryos (Martyn et al. 2018) or 3D cellular models derived from hESC (Simunovic et al. 2019; Moris et al. 2020). While these powerful approaches provide valuable insights, currently there is no *in vivo* data on the molecular control of human gastrulation to compare them against or further refine them.

Here we present a morphological and spatially resolved single cell transcriptomic characterisation of human gastrulation at Carnegie Stage (CS) 7, equivalent roughly to 16 –19 days post-fertilization, providing a detailed description of cell types present at this previously unexplored and fundamental stage of human embryonic development. We find that primordial germ cells and relatively mature blood progenitors are already present at this early embryonic stage, providing a novel perspective into the progression of cell type specification in humans. The information on the spatial origin of cells, in addition to aiding cell type identification, provides insight into the transcriptional profile of cell types in distinct anatomical regions such as the differential patterns of expression in mesoderm collected from caudal and rostral portions of the embryo. Moreover, while many aspects of gastrulation are similar between the human and mouse, we find several important differences as well. Molecules that play a key role in gastrulation and patterning in the mouse are not detected in the human, indicative of specific mechanistic differences. Finally, this transcriptomic resource provides the first transcriptional definition of the *in vivo* primed pluripotent state and serves as a refence against which *in vitro* model systems can be assessed.

## RESULTS

### Morphological and transcriptional characterization of a CS7 human gastrula

Through the Human Developmental Biology Resource (HDBR, http://www.hdbr.org; Methods) we obtained an exceptional gastrulation stage human embryo, from a donor who generously provided informed consent for the use in research of embryonic material arising from the termination of her pregnancy. The embryo was karyotypically normal and male (Region specific assay: (13, 15, 16, 18, 21, 22) x 2, (X, Y) x 1).

The sample was completely intact and morphologically normal, comprising an embryonic disk with amniotic cavity, connecting stalk and yolk sac with pigmented cells (Figure 1a). We manually micro-dissected away the yolk sac and connecting stalk to isolate the oval embryonic disk with overlying amnion. A dorsal view of the disk showed the primitive streak (PS) extending approximately half the diameter of the disk along the long, rostral-caudal, axis. The early primitive node was visible at the rostral end of the streak (Figure 1b). A ventral view showed the PS, node and forming prechordal plate (Figure 1c). The length of the primitive streak relative to the embryonic disk, presence of prechordal plate and the node at the middle of the disk allowed us to stage the embryo as Carnegie Stage (CS) 7 (O‗Rahilly and Müller 1987). This is roughly equivalent to a late-streak stage mouse embryo at embryonic day (E) 6.75–7.5 (Theiler stage 10b (Lawson and Wilson 2016)).

**Figure 1:**
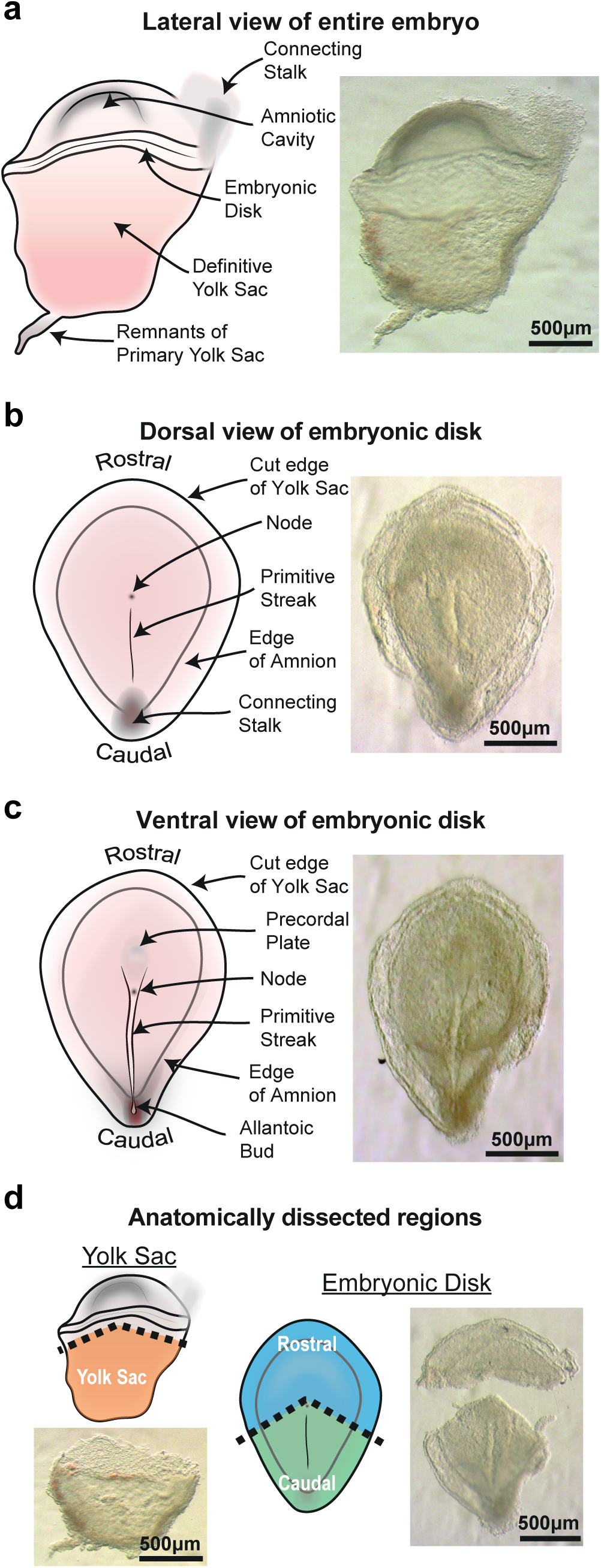
Morphological characterization of a CS7 human gastrula. a, Schematic and bright field images of an intact CS7 human embryo highlighting key morphological structures from a lateral view. Dorsal (b) and ventral (c) views of the dissected embryonic disk showing the primitive streak, node and forming bud of the allantois. d, Brightfield images showing embryo dissection with schematic diagrams highlighting the three anatomical regions collected (yolk sac, rostral and caudal regions of embryonic disk)

We used this sample to perform a spatially regionalised single-cell RNA-seq (scRNAseq) analysis. To retain anatomical information when disaggregating cells for the scRNAseq analysis, we further subdissected the embryonic disk into rostral and caudal regions (Figure 1d). The caudal region incorporated the entire PS from the node to the allantoic bud. We separately dissociated the rostral and caudal embryonic disk and the yolk sac to single cells and used FACS in combination with a live/dead stain to collect live cells for scRNAseq.

As a user-friendly community resource, we have created a web-interface to interrogate these data, accessible at http://www.human-gastrula.net.

In total, after stringent quality filtering, we generated a library of 1,195 single cells (665 caudal derived cells, 340 rostral derived cells and 190 Yolk sac derived cells) using the Smart-Seq2 protocol (Picelli et al. 2014) and Illumina sequencing, with a median of 4000 genes detected per cell (Supplementary Figure 1). We could distinguish any maternal cells from male embryonic cells on the basis of their transcriptome. All cells showed expression of Y-chromosome genes and XIST transcript was largely undetectable (Supplementary Figure 2a), confirming that there was no maternal cell contamination.

An analysis of cell cycle stage of sequenced cells revealed that all stages, including G1, G2/M and S phase could be detected, suggesting that normal cell cycling was occurring (Supplementary Figure 2b). We used the transcriptomic data to also infer the genomic integrity of the sample by estimating the number and the size of insertions and deletions (indels). This showed that in comparison with single cell transcriptomes from human fetal liver (Segal et al. 2019), the cells from our sample fall within the normal range for indels (Supplementary Figure 2c). These analyses, alongside the karyotyping (see above) and the intact morphology of the sample (Figure 1), suggest that it is representative of normal human gastrulation.

After identifying genes with highly variable expression, we detected 11 different cell populations with unsupervised clustering (Figure 2a). Using a combination of anatomical location and known marker genes, we annotated the 11 clusters as: Epiblast, Ectoderm, Primitive Streak, Nascent Mesoderm, Axial Mesoderm, Emergent Mesoderm, Advanced Mesoderm, Yolk Sac Mesoderm, Endoderm, Hemogenic Endothelial Progenitors and Erythrocytes (Figure 2a, 2b and Supplementary Table 1). The pluripotent Epiblast could be detected by the expression of *SOX2, OTX2, CDH1* and was represented in both the caudal and rostral regions of the embryo (55% caudal, 45% rostral, 0% yolk sac; Figure 2c and Supplementary Table 2). In contrast, the Ectoderm came predominantly from the rostral portion of the embryo and did not express pluripotency markers but had high expression of key markers such as *DLX5, TFAP2A* and *GATA3*, representing embryonic and amniotic ectoderm (Yang et al. 1998; Streit 2007). The Primitive Streak, was identified by the archetypal marker *TBXT* (*Brachyury*) in combination with *CDH1* and *FST* (Figure 2b, e). As expected, these cells originated almost exclusively from the caudal portion of the embryo. Whilst the majority of Nascent Mesoderm cells were also located in the caudal region and expressed *TBXT*, they could be distinguished from PS cells by the expression of key mesodermal markers *MESP1* and *PDGFRA* (Figure 2b, e). This co-expression of both PS and mesoderm markers led us to define this mesoderm as ‘nascent’, representing the forming mesoderm cells in the process of delaminating from the PS. Axial Mesoderm, which gives rise to the notochord, the midline rod-like structure that is the defining feature of the chordate phylum to which humans belong, could be detected by the expression of *TBXT, CHRD* and *NOTO*.

**Figure 2:**
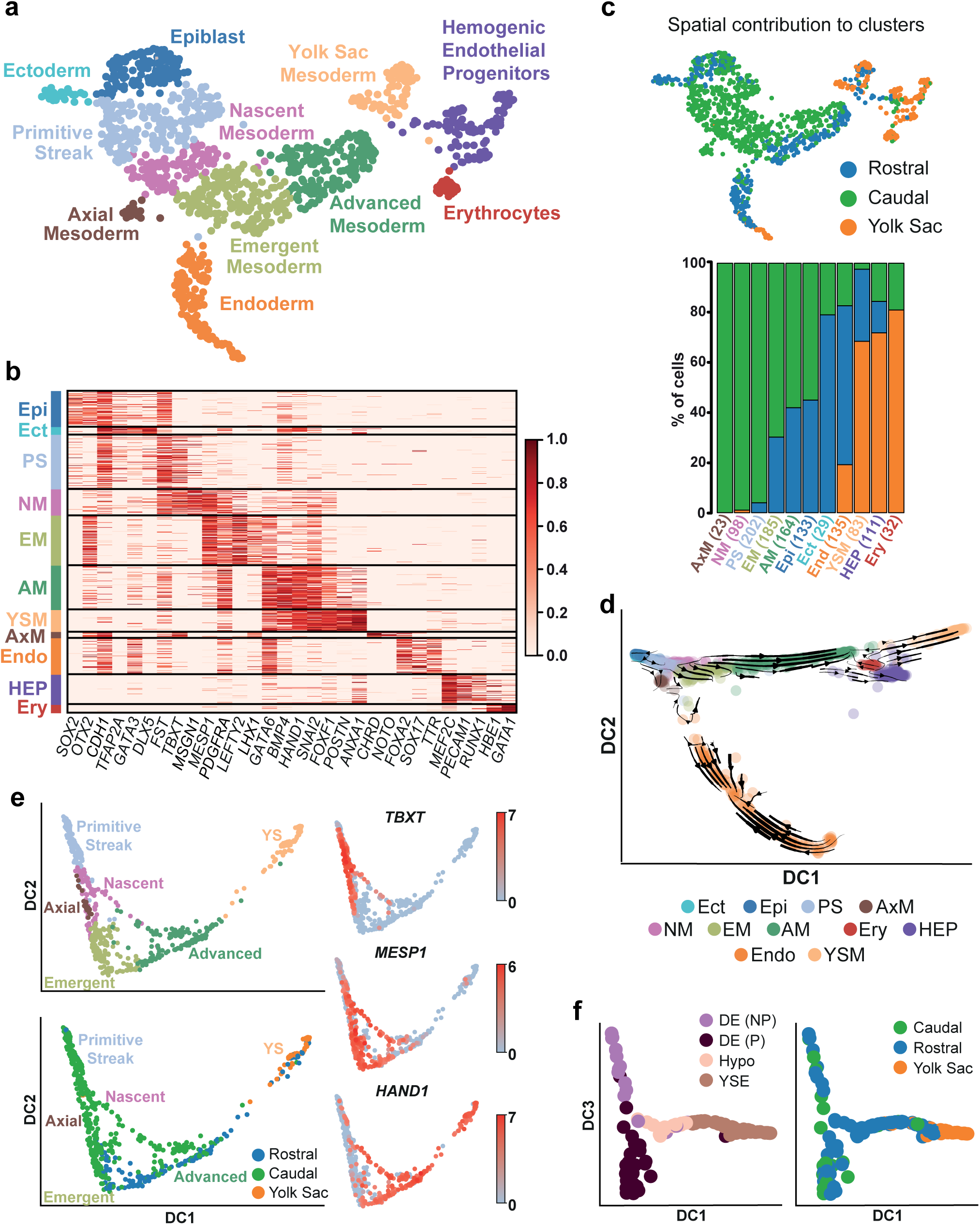
Spatially resolved single cell transcriptomic analysis of a gastrulating human embryo. a, UMAP plot of all the cells that passed quality control (n = 1,195) computed from highly variable genes. Cells with similar transcriptional profiles were clustered into 11 different groups, as indicated by the different colours. b, Heatmap with the expression of well characterized marker genes for the identified cell types: Epiblast (Epi), Ectoderm (Ect), Primitive Streak (PS), Nascent Mesoderm (NM), Emergent Mesoderm (EM), Advanced Mesoderm (AM), Yolk Sac Mesoderm (YSM), Axial Mesoderm (AxM), Endoderm (Endo), Hemogenic Endothelial Progenitors (HEP), Erythrocytes (Ery). Each gene’s normalized log expression levels are standardized so that they vary within [0,1]. c, UMAP and bar plot highlighting the anatomical region that cells were collected from and the percentage breakdown of each cluster. Numbers in brackets represent the total number of cells per cluster. d, RNA velocity vectors overlaid on diffusion map of cells from all 11 clusters; the first two diffusion components (DC1 and DC2) are shown. e, Diffusion map of cells from the 6 mesoderm related clusters (Primitive Streak, Nascent Mesoderm, Emergent Mesoderm, Advanced Mesoderm, Axial Mesoderm and Yolk Sac (YS) Mesoderm); the first two diffusion components (DC1 and DC2) are shown. In the top left panel, cluster identity is indicated by the different colours. The bottom left panel shows the anatomical location that cells were collected from. The right panels show the expression of key mesodermal marker genes highlighting the heterogeneity in mesoderm types. f, Diffusion map of endodermal cells showing diffusion components (DC) 1 and 3. In the diffusion map on the left endodermal sub clusters are marked by different colours, while on the right the anatomical location from which cells were collected is shown.

Two other clusters of embryonic mesoderm could be distinguished by their relative degree of maturation and location within the embryo. We annotated the first as Emergent Mesoderm since it expressed the highest levels of *MESP1* but was negative for *TBXT*, thereby representing a transition from the Nascent Mesoderm towards the more mature Advanced Mesoderm (Figure 2e). It also expressed *LHX1* and *OTX2* as well as the highest levels of *LEFTY2*, which in the mouse is expressed in mesoderm arising from the mid-distal region of the PS ((Meno et al. 1999). The Advanced Mesoderm cluster was relatively more mature based on the decreased expression of *MESP1* and the highest expression of mesoderm marker genes *PDGFRA* and *GATA6*. The Advanced Mesoderm cluster also expressed *HAND1, BMP4, FOXF1* and *SNAI2*, all of which are markers of relatively more mature mesoderm (Figure 2b, e).

The information on the spatial origin of the cells allowed us to track the progression of mesoderm maturity. Cells that had more time (since they emerged from the PS) to mature might also, in that time, be expected to have migrated further from the PS. Consistent with this, Nascent Mesoderm was almost entirely collected from the caudal portion of the embryo (99% caudal, 0% rostral, 1% yolk sac; Figure 2c and Supplementary Table 2). Similarly, Axial Mesoderm was only located in the caudal region, consistent with it having just emerged from the PS when the embryo was collected. In contrast, Emergent Mesoderm and Advanced Mesoderm were collected from both the rostral and caudal regions of the embryo (Emergent: 70% caudal, 30% rostral; Advanced: 58% caudal, 42% rostral), highlighting that they had migrated rostrally away from the PS. These two mesoderm clusters also showed evidence of sub-structure based on Rostral-Caudal differences in origin (Figure 2e).

Yolk Sac Mesoderm was identified based both on its anatomical origin (69% yolk sac, 29% rostral, 2% caudal) as well as the expression of specific marker genes such as *POSTN* and *ANXA1*. This cluster showed overlapping expression of markers such as *HAND1, FOXF1* and *SNAI2* with the Advanced Mesoderm. Other mesoderm-derived clusters included Erythrocytes, which had high expression of hemoglobin genes including hemoglobin *HBE1* as well as the blood-related transcription factor *GATA1*. The majority of erythrocytes were collected from the yolk sac (81% yolk sac, 19% caudal, 0% rostral). We annotated a Hemogenic Endothelial Progenitor population based on the expression of both endothelial makers (*PECAM1* and *MEF2C*) as well as hematopoietic markers (*RUNX1* and *GATA1*). Hemogenic Endothelial Progenitor cells were also located predominantly in the yolk sac, although some cells also came from caudal and rostral regions (72% yolk sac, 15% caudal, 12% rostral).

Endoderm could be identified by the expression of *SOX17, GATA6, FOXA2* and *TTR*. Endoderm was collected from all three anatomical regions (64% rostral, 19% yolk sac, 17% caudal; Figure 2c). Further sub-clustering of the Endoderm revealed the different endodermal subtypes, including two populations of PS derived Definitive Endoderm, primitive endoderm derived Hypoblast and extra-embryonic Yolk Sac Endoderm (Figure 2f and Supplementary Figure 3).

To understand how cells transition from pluripotent epiblast to the different germ layers, we ordered cells along differentiation trajectories using diffusion maps and RNA velocity analysis (Haghverdi et al. 2016; La Manno et al. 2018) (Figure 2d and Supplementary Figure 4). This revealed trajectories from the Epiblast population along three broad streams, corresponding to the three germ layers mesoderm, endoderm and ectoderm. The first two are separated mainly along the first and the second diffusion components (DC1 and DC2; Figure 2d), while Ectoderm is present as a small separate cluster along the third diffusion component (DC3; Supplementary Figure 4). Because it corresponds closely to the cell types and their spatial location, DC1 reflects the extent of their differentiation and the ‘age’ of cells, based on how far in the past of this sample they emerged from the Epiblast (Figure 2d and Supplementary Figure 4). For instance, cell types such as extra-embryonic mesoderm which emerge relatively early during gastrulation are plotted further from the epiblast than cell types such as axial mesoderm that emerge later. The extra-embryonic mesoderm in humans, in addition to being formed by the earliest cells to emerge from the streak, is understood to also have contribution from the hypoblast of the bilaminar disk stage embryo (Bianchi et al. 1993). The high DC1 value of these cells is consistent with their early origin prior to and during gastrulation.

### Testing in vitro models of human development

Due to the difficulty of accessing early human embryos, there is an increasing effort to produce *in vitro* models of human development, particularly during gastrulation (Moris et al. 2020). Testing the extent to which these *in vitro* models accurately represent the *in vivo* state is crucial and the comprehensive characterization of cell types transcriptome and spatial location in a human CS7 embryo presented here offers a unique opportunity to do this.

We used our data to test rostral-caudal patterning in hESC derived gastruloids (Moris et al. 2020). Spatial transcriptomic data from human gastruloids provides gene clusters that have been ordered according to their pattern of expression along the rostral-caudal axis. We exploited our data to identify genes that are characteristic of cell populations that are enriched in the caudal or the rostral region of the embryo (see Methods) and compared them with the gastruloid gene clusters. By doing so, we could validate the broad rostral-caudal pattern present in gastruloids (Supplementary Figure 5a). Moreover, we also revealed which gastruloids gene clusters most closely correspond to human cell clusters, information that could be used in future to further refine protocols for achieving precise patterning of *in vitro* models (see Methods and Supplementary Figure 5a).

Another important effort is towards the creation of accurate *in vitro* models of human pluripotency. *In vitro*, the naïve as well as the primed pluripotency states have been characterized in human embryonic stem cells (hESC) (Messmer et al. 2019). However, *in vivo* the transcriptional signature of only naïve pluripotency has been captured from sequencing pre-implantation human embryos, and it is not clear how long the naïve state persists in the developing human epiblast. The CS7 Epiblast cluster offered the opportunity to transcriptionally characterize for the first time the human *in vivo* primed pluripotent state, and thereby define the transition from naïve to primed pluripotency. To this end, we compared the transcriptomes of pluripotent cells from pre-implantation human embryos (Petropoulos et al. 2016) against the CS7 Epiblast using a Principal Component Analysis (PCA) (Figure 3a). Cells showed an ordered pattern according to their developmental stage along PC1 and PC2, showing that the CS7 Epiblast is transcriptionally distinct from even the E7 epiblast. Having anchors for the *in vivo* naïve and primed states allowed us to project the transcriptomes of naïve and primed hESC onto this PCA. We found that naïve hESC plotted closest to E6/E7 cells while primed hESC plotted closest to and partially overlapped with CS7 Epiblast. Indeed, naïve hESC showed the highest correlation with the transcriptome of E6 epiblast, while primed hESC correlated most closely with CS7 Epiblast (Supplementary Figure 5b). This verifies that the primed state captured *in vitro* in hESC does closely represent at the global transcriptome level the *in vivo* primed state.

**Figure 3:**
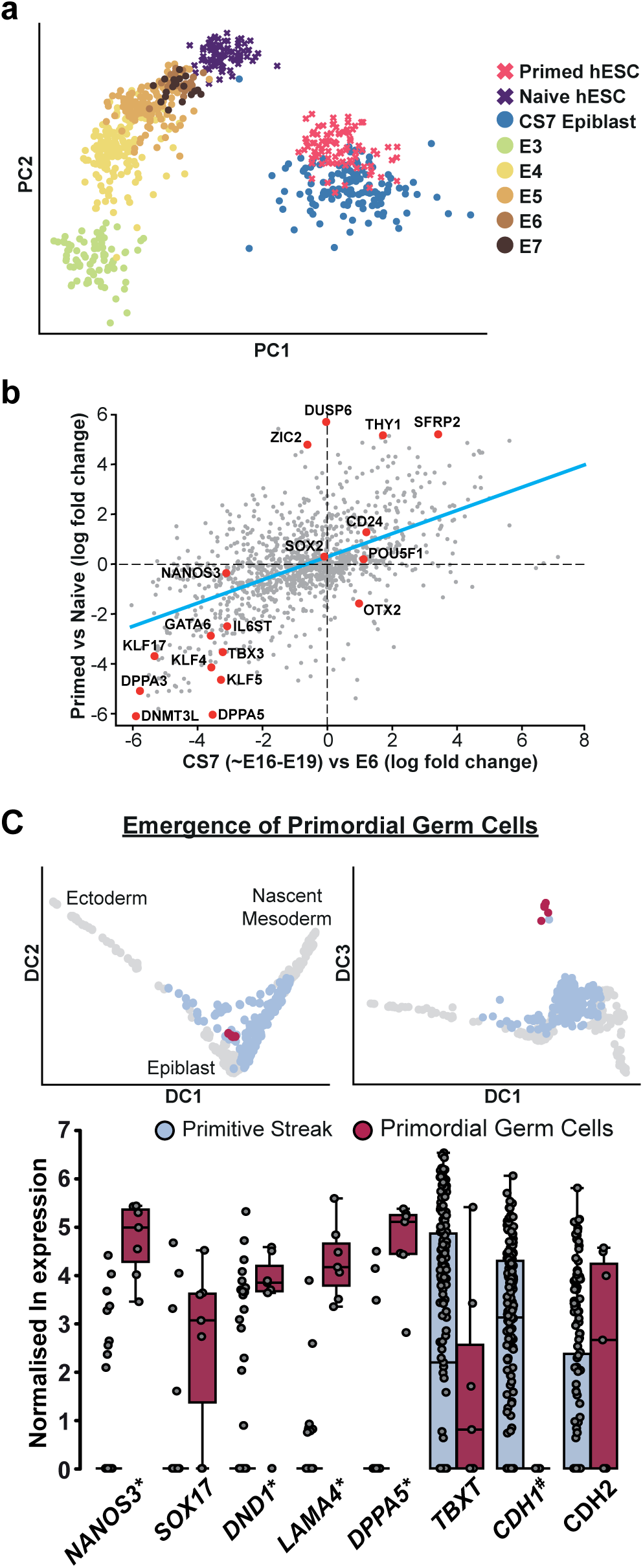
Comparison of an in vitro model of pluripotency with in vivo data and identification of primordial germ cells. a, Principal component analysis (PCA) of the transcriptomic profiles of pluripotent cells from pre-implantation human embryos (between embryonic day (E) 3 and 7; data from (Petropoulos et al. 2016)) and of the epiblast cells from the CS7 human gastrula. Single-cell RNA-seq data from hESC in a primed and naïve state (data from (Messmer et al. 2019)) were projected on top of this PCA. b, Log-fold changes of expression levels of the genes used for the PCA between primed vs naïve hESC (y axis) and CS7 epiblast vs E6 data (x axis). Selected genes are highlighted in red; the blue line is obtained through a linear regression. A strong, positive correlation is found (Pearson’s correlation coefficient ∼0.63), indicating that the hESC resemble the *in vivo* primed and naïve states at the transcriptome-wide level. c, Unbiased search for rare cell types with RACEID (see Methods) revealed the emergence of a Primordial Germ Cell (PGC) population in the Primitive Streak cluster. PGCs are shown in red on a diffusion map including Epiblast, Primitive Streak, Ectoderm and Nascent Mesoderm. Diffusion component (DC) 3 highlights a clear separation between the PGCs and the Primitive Streak (PS). The boxplot shows the normalized log expression of key PGC and PS marker genes in the PGC sub-cluster compared to the PS cluster. The stars and hashes mark the names of genes that are statistically significantly higher or lower respectively in PGCs compared to PS.

Another confirmation of this was obtained by comparing the log-fold changes of genes between E6 vs CS7 epiblast against primed vs naïve hESC (Figure 3b), which strongly correlated (Pearson’s correlation coefficient ∼0.63). In particular, this analysis revealed compatible trends of previously known markers of the naïve (eg, *KLF5, KLF17, DNMT3*) as well as of the primed state (eg, *CD24, THY1, SFRP2*; Figure 3b) (Messmer et al. 2019). However, there are several genes that showed opposite behaviors *in vivo* and *in vitro*, providing targets for manipulations that could achieve a ‘truer’ primed state *in vitro*.

### Primordial Germ Cells can be detected as early as CS 7 in vivo

An important population of cells to originate from the early Epiblast are the Primordial Germ Cells (PGCs), the highly specialised precursor cell type that bridge the generations by giving rise to the gametes – ova in females and sperm in males. In the mouse, PGCs emerge at around E7.25 (Chiquoine 1954; Magnúsdóttir and Azim Surani 2014). Recent work has shown that cells expressing some PGC markers can be identified at E11 (Sasaki et al. 2016) in non-human primates and in *ex vivo* cultured human embryos (Chen et al. 2019). It is unknown when PGCs start to develop during human development *in utero*.

In humans, *SOX17* is a critical specifier of PGC fate (Irie et al. 2015). In our data, *SOX17* was mainly expressed in the endoderm. However, we detected a limited number of SOX17 expressing cells in the PS population that might represent PGCs. To test this possibility, we used an unbiased clustering algorithm, RACEID, to identify rare cell populations within our PS cluster. This revealed a sub-cluster of cells that represent PGCs as they expressed low levels of *TBXT* and *CDH1* and high levels of several well characterized PGC markers including *NANOS3, SOX17, DND1, DPPA5* and *LAMA4* (Magnúsdóttir and Azim Surani 2014) (Figure 3c and Supplementary Table 3 for a list of genes up- or down-regulated in PGCs). Notably, they also clustered as a distinct group on the diffusion map of PS, Ectoderm, Epiblast and Nascent Mesoderm (Figure 3c). Consistent with recent *in vitro* data (Chen et al. 2019) this suggests that by CS7, the human embryo has already started to set aside PGCs.

### Early differentiation of the Epiblast

To probe in more detail the changes triggered in the epiblast by gastrulation, we plotted a diffusion map with the sub-set of cells belonging to the Epiblast, Ectoderm, Primitive Streak and Nascent Mesoderm clusters (Figure 4). We also computed RNA velocity vectors (La Manno et al. 2018) and overlaid them onto the diffusion map, to provide added information about the differentiation trajectories of these cells. This analysis supported the existence of a bifurcation from Epiblast, on one side towards Mesoderm via the Primitive Streak and on the other side towards Ectoderm (Figure 4a). This was also reflected in the anatomical origin of cells (Supplementary Figure 6a).

**Figure 4:**
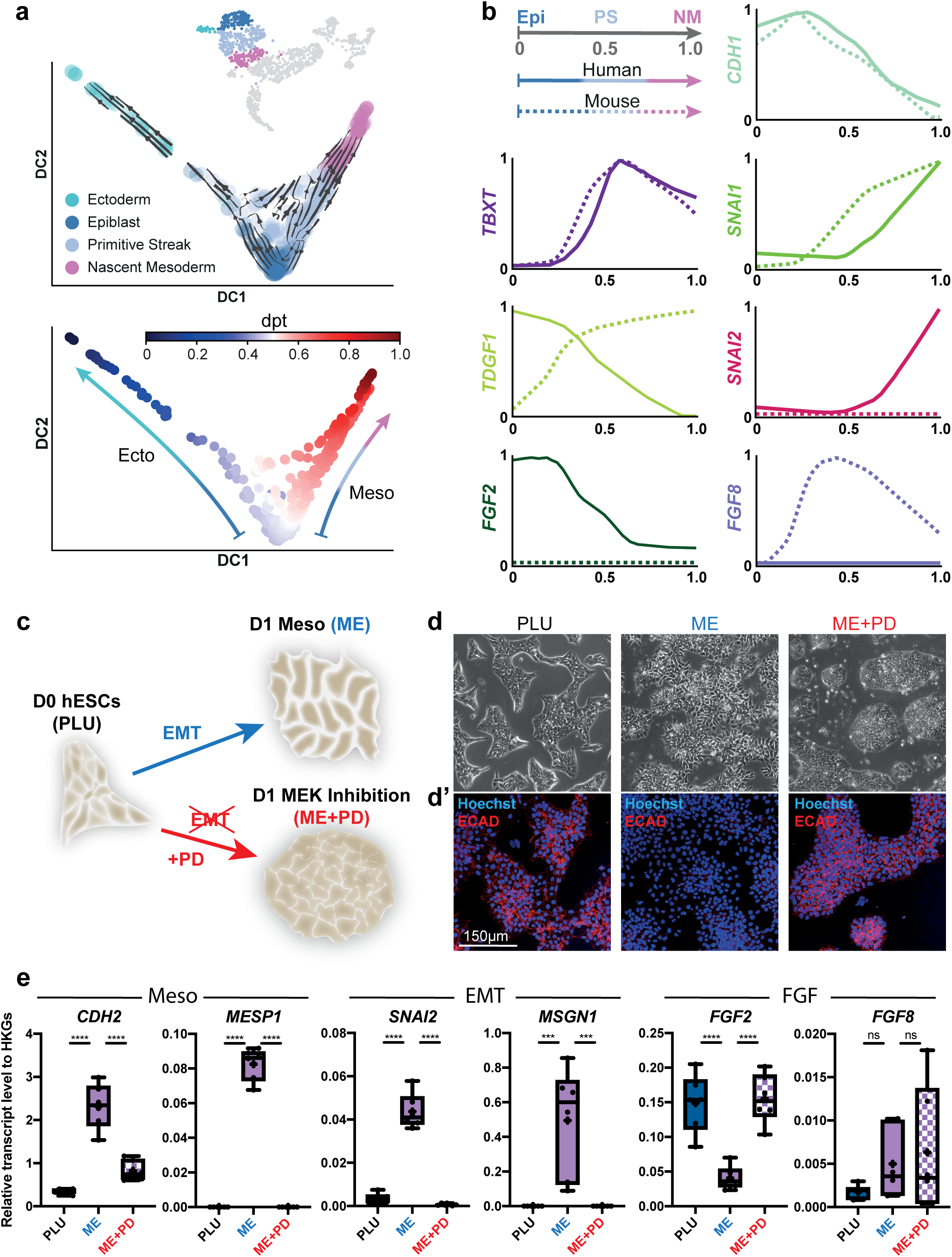
Characterization of EMT during human and mouse gastrulation. a, Transcriptional changes during epiblast differentiation. The top panel shows a diffusion map and RNA velocity vectors of 4 clusters: Epiblast, Primitive Streak, Nascent Mesoderm and Ectoderm (highlighted in the inserted UMAP plot). The first two diffusion components are shown (DC1 and DC2). The bottom diffusion map shows the diffusion pseudotime (dpt) coordinate, with the arrows on either side indicating the direction of the two differentiation trajectories originating from Epiblast (with dpt ∼ 0.5) and moving towards Ectoderm (dpt ∼ 0) or Mesoderm (dpt ∼ 1). b, Comparison of pseudotime analysis during primitive streak and nascent mesoderm formation in human and mouse (data from (Pijuan-Sala et al. 2019)). Cells in epiblast (Epi), Primitive Streak (PS) and Nascent Mesoderm (NM) clusters from human and mouse embryos at matching stages (see Methods) were independently aligned along a differentiation trajectory and a diffusion pseudotime coordinate (dpt) was calculated for each of them (top left). The expression pattern of selected genes along pseudotime is plotted for human (continuous lines) and mouse (dashed lines). Whilst CDH1, TBXT and SNAI1 followed the same trends between species, differences could be observed in SNAI2, TDGF1, FGF8, FGF2. c, Schematic representation of an in vitro model for epithelial-to-mesenchymal transition (EMT) and its inhibition. Human pluripotent embryonic stem cells (D0 hESC, PLU) are differentiated towards Mesendoderm (D1 MESO, ME) and undergo epithelial-to-mesenchymal transition (EMT). However, if MEK pathway is inhibited, EMT is prevented (D1 MEK Inhibition, ME+PD). d, d’ Bright-field (top) and fluorescence microscopy images of different ESC colonies(bottom; Hoechst staining and immunofluorescence for ECAD) of D0 hESC (left), D1 Meso (center) and D1 MEK Inhibition (right). e, Quantification of transcript levels for selected genes across the three conditions PLU, ME, ME+PD. Y-axis represents expression levels relative to house keeping genes (HKGs). Mesodermal (CDH2 and MESP1) and EMT (SNAI2 and MSGN1) markers increase from PLU to ME, but their increase is abolished upon MEK inhibition (ME+PD). Conversely, FGF2 is downregulated during EMT, while it stays up if EMT is blocked. FGF8 does not show any statistically significant changes across these conditions. qPCR results confirm our predictions from in vivo data (panel b). (n = 6 from three different experiments. ns = p-value ≥ 0.05; *** = p-value < 0.001; **** = p-value < 0.0001; ordinary one-way ANOVA after Shapiro-Wilk normality test).

Ordering cells using diffusion pseudotime provided a method to infer the changes in gene expression as Epiblast cells enter the Primitive Streak and begin to delaminate to form the Nascent Mesoderm (Figure 4a and Supplementary Figure 6). Epiblast markers such as *SOX2* and *OTX2* decreased as cells differentiated towards Nascent Mesoderm. The expression of the pluripotency marker *POU5F1* (*OCT4*) however was maintained during transition towards Nascent Mesoderm (Supplementary Figure 6b). Ectodermal specification could be detected by the increased expression of *DLX5, TFAP2A* and *GATA3* (Streit 2007), representing the early specification of non-neural ectoderm, but also consistent with the extra-embryonic ectoderm of the amnion (Supplementary Figure 6c). Whilst ectoderm specific transcripts showed a robust increase, markers of early neural induction such as *SOX1, SOX3, PAX6* and markers of differentiated neurons such as *TUBB3, ELAVL3, OLIG2, NEUROG1, NEUROD1* or *NKX2*.*2* were extremely low or undetectable (Supplementary Figure 6c) (Trevers et al. 2017; Delile et al. 2019). Together, these data show that there are no neural ectoderm cells in the CS7 embryo.

### Signaling differences during human and mouse gastrulation

Epithelial cells of the epiblast undergo an epithelial to mesenchymal transformation (EMT) by downregulating adherens junction molecules such as E-Cadherin (*CDH1*) so they can delaminate from the epiblast and migrate away as mesenchymal cells. In the mouse embryo, E-Cadherin expression is downregulated by the transcriptional repressor Snail (Cano et al. 2000), and FGF signaling is essential for the migration of delaminated cells (Sun et al. 1999).

Pseudotime analysis showed that in the CS7 gastrula, the PS marker *TBXT* (*Brachyury*) increased during the transition from Epiblast to Nascent Mesoderm, peaking in the Primitive Streak, while as expected, the mesoderm marker *MESP1* increased with the formation of Nascent Mesoderm (Figure 4b). A core event during EMT in mouse is a switch in the adherens junction molecules Cadherins, from E-to N-Cadherin (*CDH1* to *CDH2*) (Smith et al. 1992; Cano et al. 2000). In the human, we could detect a similar trend in these genes, including the Cadherin switch, with *CDH1* decreasing towards Nascent Mesoderm while *CDH2* increased (Figure 4b and Supplementary Figure 6b). In addition, we found 3,350 genes that were differentially expressed along the developmental trajectory between Epiblast and Nascent Mesoderm (see Supplementary Table 4).

While we observed several broad similarities in the formation of mesoderm between the human and other model organisms, we wanted to test this in more detail in an unbiased manner. To do this, we used pseudotime analyses to compare the transition from epiblast to early mesoderm in the human cells with the equivalent populations in the Mouse Gastrula Single Cell Atlas (Pijuan-Sala et al. 2019) (Supplementary Figure 7). We identified 662 genes that were differentially expressed along the developmental trajectories from Epiblast to Nascent Mesoderm in either human or mouse (Supplementary Figure 7 and Supplementary Table 5). Of these, 531 genes shared the same trend across pseudotime, either increasing (117) or decreasing (414). For example, in both mouse and human, *CDH1* decreased during transition from epiblast to nascent mesoderm, *TBXT* increased before starting to decrease and *SNAI1* continuously increased towards nascent mesoderm (Figure 4b). However, there were some notable differences.

One example was in the expression of the zinc-finger transcription factor *SNAI2* (Slug), a regulator of EMT. *SNAI2* levels increased dramatically during Nascent Mesoderm formation in the human. However, *SNAI2* was not detected during this transition in the mouse (Figure 4b), as also confirmed by additional independent mouse transcriptomic datasets from the same stages (Peng et al. 2016). The lack of requirement for *SNAI2* during mouse gastrulation is consistent with the viability of *SNAI2* null mice (Jiang et al. 1998). By contrast, in the chick as in the human, *SNAI2* is expressed within the PS and interfering in its expression resulted in the impaired emergence of mesoderm from the PS (Nieto et al. 1994). Together this suggests that unlike in the mouse, in human, *SNAI2* may play a role in regulating EMT during gastrulation.

In the mouse, the expression of various signaling molecules is crucial for EMT, germ layer specification and migration (Ciruna and Rossant 2001; Ding et al. 1998; Jin and Ding 2013). Our analyses again revealed striking differences between the mouse and human in the expression and trends of these signals. In the mouse, *TDGF1*, a NODAL co-receptor essential for normal mesodermal patterning, shows an increase in expression during Primitive Streak and Nascent Mesoderm formation. In contrast, in the human gastrula *TDGF1* expression showed the opposite trend, decreasing as Nascent Mesoderm formed (Figure 4b). *FGF8* is the only known FGF directly required for gastrulation (Ornitz and Itoh 2015) in the mouse, playing a particularly important role in the migration of cells away from the PS (Sun et al. 1999). By contrast, FGF8 was completely absent during the transition from Epiblast to Nascent Mesoderm in human (Figure 4b). Other FGF members are expressed during this transition however, including *FGF4* (which is also expressed in the mouse), and *FGF2*, which is not required for gastrulation in the mouse (Zhou et al. 1998; Ortega et al. 1998) nor expressed, as confirmed in other datasets (Peng et al. 2016) (Figure 4b). In human, unlike for *FGF8*, there are no reported diseases associated with *FGF2* mutations, suggesting that mutations in *FGF2* may not be viable (Ornitz and Itoh 2015). Interestingly treatment of in vitro cultured mouse epiblasts with FGF2 resulted in the altered fate of these cells from ectoderm to mesoderm (Burdsal et al. 1998), pointing to a degree of redundancy in function between these FGFs.

To experimentally validate these changes in expression of key molecules during the EMT accompanying the transition from pluripotency to mesoderm, we used a hESC based differentiation model. We cultured hESC under conditions that promote differentiation (Teo et al. 2011; Mendjan et al. 2014; Yiangou et al. 2019) towards a mesendoderm progenitor state (ME), which corresponds to Primitive Streak and Nascent Mesoderm states in Figure.2. During this specification, hESC colonies showed hallmarks of the EMT associated with gastrulation, such as a dispersed morphology and downregulation of E-Cadherin (Figure 4c, d). This EMT can be blocked using PD0325901, an inhibitor of MAPK/ERK kinase (MEK), that is the major downstream pathway of FGF signaling (Thisse and Thisse 2005) (Figure 4c, d, d’). We next used this system to test gene expression changes during gastrulation specific to humans. The expression of gastrulation markers *CDH2* (N-Cadherin), *MESP1, TBXT* and *SNAI1* all increased significantly in ME as expected, and this increase could be blocked with the MEK inhibitor (Figure 4e and Supplementary Figure 8).

We found that the human-specific expression patterns that emerged from our computational analyses were also confirmed in this *in vitro* model of human gastrulation EMT. *FGF8* was expressed at very low levels and did not show any significant response to the differentiation conditions. *SNAI2* and *MSGN1* however, showed a significant increase, while *FGF2* and *TDGF1* showed a significant decrease (Figure 4e and Supplementary Figure 8). Upon MEK inhibition, all these genes were restored to their baseline values in ESC colonies, except for *TDGF1*, which suggests that this gene might be differentially regulated by multiple pathways.

Together, these results indicate that there is broad conservation of several molecular players in human and mouse gastrulation, such as the involvement of the SNAIL/SLUG family of transcriptional repressors and the influence of FGF/ MEK-dependent EMT. However, the specific members of these families vary between humans and mice. In addition, some molecules such as MSGN1, that play no role during gastrulation in mice, might do so in human. MSGN1 null mouse embryos gastrulate normally (Jeong Kyo Yoon and Wold 2000; Nowotschin et al. 2012), but show defects in regulating EMT slightly later, during somitogenesis (Chalamalasetty et al. 2014). Its robust expression during gastrulation in the human in the Nascent Mesoderm (Figure 2b) raises the possibility that its role in regulating EMT might be deployed earlier (and in a different setting) in the human compared to mouse. Overall, these results show that the comparison of our *in vivo* data with *in vitro* models for EMT in combination with inhibitors might help disentangle the roles of different pathways in the expression changes of individual genes during EMT.

### Early maturation of human hemogenic progenitors

Our initial analysis revealed two blood related clusters, Erythrocytes and Hemogenic/ Endothelial Progenitors (HEP). The identification of erythrocytes also corresponded with the observation of pigmented cells in the yolk sac and the expression of hemoglobin genes (Figure 5a and Supplementary Figure 9b). This observation was striking given the absence of pigmented blood cells at the equivalent stage in mouse embryos (∼E7.25). The expression of XIST and Y-chromosome specific genes (Figure 5a and Supplementary Figure 2a) ruled out the possibility of maternal origin of these cells.

**Figure 5:**
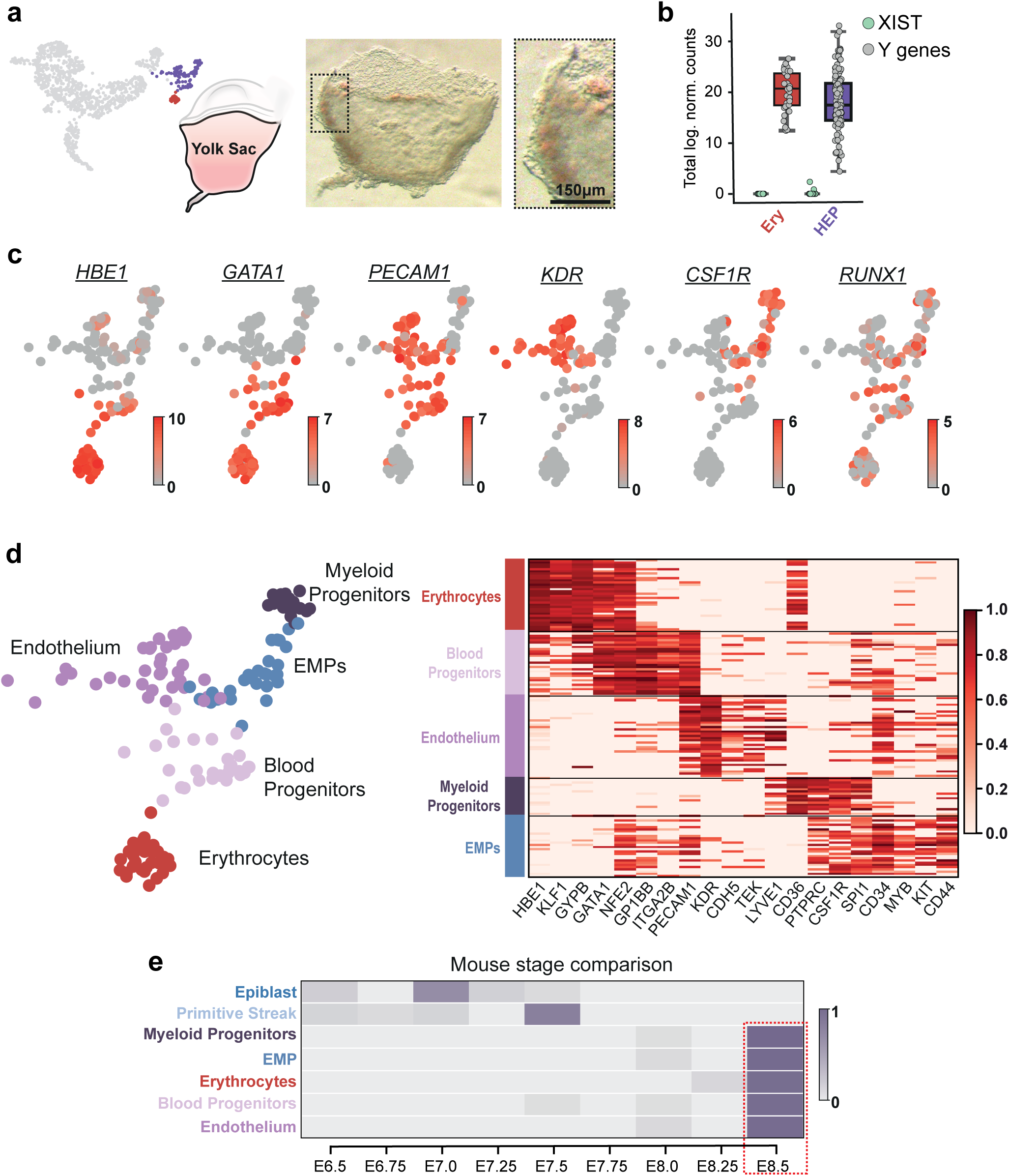
Identification of early blood progenitor types in the human. a, Brightfield image of the Yolk Sac highlighting the presence of pigmented cells. The dotted box represents the zoomed in region at right. b, Boxplots showing the total log expression of normalized counts for XIST and Y-genes in Erythrocytes (Ery) and Hemogenic Endothelial Progenitors (HEP), indicating no contamination from maternal tissue. c, UMAP plots of HEP and Erythrocyte clusters showing normalized log expression of blood related marker genes. d, UMAP plot of the HEP and Erythrocyte clusters showing four sub clusters within the HEPs, marked by the different colours. The heatmap displays the expression of well-characterized marker genes. Each gene’s normalized log expression levels are standardized so that they are within [0,1]. e, Estimation of equivalent mouse stage for selected human gastrula clusters. The heatmap includes the fraction of human cells from each cluster (rows) that maps onto analogous mouse cell types at the different stages (columns). While the majority of Epiblast and Primitive Streak cells are most similar to their mouse counterpart at E7.0 and E7.5 respectively, blood clusters are all equivalent mostly to E8.5 (dashed red rectangle).

Both blood-related populations expressed erythrocyte markers (McGrath et al. 2015); however the HEP population had a mixed expression profile of endothelial, myeloid and erythrocyte markers, suggesting a higher order substructure (Figure 5b). Unsupervised clustering of the HEP population revealed four different subpopulations with distinct transcriptional signatures (Figure 5c, Supplementary Figure 9b and Supplementary Table 6). One subpopulation represented Endothelium (Endo) based on the high expression of *PECAM1, CDH5, KDR* and *TEK*. Another expressed both megakaryocyte (*GP1BB, ITGA2B* (CD41), *NFE2*) and erythroid (*GATA1, KLF1, GYPB, HBE1*) markers, which we annotated as Blood Progenitors. A Myeloid Progenitor sub-population could be identified on the basis of high levels of monocyte/macrophage markers *CD36, CSF1R*, and *LYVE1*.

The final subcluster had an unusual transcriptional profile given the early stage of the sample, expressing a range of myeloid and erythroid markers including *KIT, CSF1R, MYB, SPI1* (PU.1), *CD34, PTPRC* (CD45), *CD52* and *NFE2* (Palis 2016). Based on these markers and the expression of *MYB*, which in the mouse marks Erythroid-Myeloid Progenitors (EMP) that constitute the second wave of macrophage progenitor at ∼E8.25, we annotated this cluster as EMP. Supporting this annotation is their co-expression of *CD34, CD45* and *CD44*, which have recently been used to define a yolk sac-derived myeloid-biased progenitor in CS11 human embryos (Bian et al. 2020). These CS11 cells have been shown to have multi-lineage potential and are thought to correspond with mouse definitive EMPs. Our results therefore indicate that such progenitors might already start to emerge as early as CS7.

Our identification of pigmented cells and multiple hematopoietic progenitor populations suggest that hematopoiesis in the human CS7 embryo has progressed further in comparison to an equivalent stage mouse embryo (E6.75–7.5). To unbiasedly examine the developmental stage of our blood related clusters in relation to the mouse, we compared the sequence of the human clusters to the equivalent populations from the Mouse Gastrula Single Cell Atlas (Pijuan-Sala et al. 2019) that spans E6.5-E8.5. Human Epiblast and Primitive Streak most closely corresponded to stages E7.0 and E7.5 respectively in the mouse, supporting our staging of the human embryo based on morphological criteria (Figure 5e). In contrast, the human blood populations all most closely correlated with developmental stage E8.5 in the mouse (Figure 5e). This suggests that hematopoiesis is further advanced in the human compared to the equivalent stage in mouse. This might be because the cellular differentiation program during hematopoiesis progresses over similar timescales in the human and mouse, but the process of gastrulation is of longer duration in the former (over a week in humans compared to <4 days in mouse), resulting in mature hematopoietic cells being present at relatively early stages of human gastrulation. Our analyses further indicate that in comparisons of the human and mouse, different cell and tissue types will not necessarily develop in synchrony with each other.

## DISCUSSION

Our characterization of human gastrulation combines single cell transcriptomics with manual microdissection to leverage the added information of the anatomical origin of cells. This reveals that the human embryo already has PGCs and red blood cells, but has not yet initiated neural specification at CS7 (∼16 to 19 days post fertilization). The differentiation trajectory of gastrulating cells from epiblast to mesoderm is broadly conserved between humans and the mouse, but specific differences in signaling molecules suggest that the process of EMT may be regulated differently in the human compared to the mouse. These human specific details of differentiation will be a valuable resource for developing approaches for the directed differentiation of human embryonic stem cells. Furthermore, they will help in interpreting experimental results on gastrulation derived from model organisms such as the mouse, or *in vitro* gastruloid systems.

The human and mouse gastrula are morphologically very different, the former being a disc and the latter being cylindrical. This profound difference in morphology alters the migratory path of cells during gastrulation and therefore the inductive signals cells might be subject to from neighbouring germ layers. It will therefore be important to compare this human gastrula single-cell transcriptome to that from other organisms with a similar embryonic disc, such as the rabbit, chick and non-human primates. This will enable us to address to what extent the specific differences between human and mouse transcriptomes are due simply to evolutionary divergence or rather, reflect difference in morphology.

This unique data-set of a complete post-implantation human embryo provides the first glimpse into a largely inaccessible stage of our development and represent a valuable resource for the design of strategies for directed differentiation of human stem cells for therapeutic and research ends.

## METHODS

### Collection of Human Gastrula Cells

The CS7 embryo was provided by the Human Developmental Biology Resource (HDBR -https://www.hdbr.org/general-information). HDBR has approval from the UK National Research Ethics Service (London Fulham Research Ethics Committee (18/LO/0822) and the Newcastle Tyneside NHS Health Authority Joint Ethics Committee (08/H0906/21+5)) to function as a Research Tissue Bank for registered projects. The HDBR is monitored by The Human Tissue Authority (HTA) for compliance with the Human Tissue Act (HTA; 2004). This work was done as part projects (#200295 and #200445) registered with the HDBR. The material was collected after appropriate informed written consent from the The sample was collected in cold L15 media and initially characterised by the HDBR. It was then transferred to M2 media and imaged on a Stereo microscope. The sample was micro-dissected using tungstenneedles and dissociated into single cells using 200μl Accutase (ThermoFisher, Cat No. A1110501) for 12 minutesat 37°C, being agitated every 2 minutes, before adding 200μl heat-inactivated FBS (ThermoFisher, Cat No. 10500) to quench the reaction. Cells were then centrifuged atfo1r030m0ripnmutes at 4°C before being suspended in 100μl HBSS (ThermoFisher, Cat No. 14025) + 1% FBS, and stored on ice. Single cells were collected using a Sony SH800 FACS machine w stringent single-cell collection protocol and sorted into 384 well plates containing SMART-seq2 lysis buffer (Picelli et al. 2014) plus ERCC spik (1:10M). To ensure we collected good quality cells, a live/dead d(yAebcam, Cat No. ab115347) was used; 100μl was added to the cell suspension at a 2x concentration in HBSS 10 minutes before collection, and live cells were collected based on their FITC intensity. Once cells were collected, plates were sealed, spun down, and frozen using dry ice before being stored at −80°C. This complete process, from dissection to single-cell collection, took approximately 2-3 hours.

### Single-cell RNA sequencing

mRNA from single cells was isolated and amplified (21 PCR cycles) using the SMART-seq2 protocol (Picelli et al. 2014). Multiplexed sequencing libraries were generated from cDNA using the Illumina Nextera XT protocol and 125 bp paired-end sequencing was performed on an Illumina HiSeq 2500 instrument (V4 chemistry).

### Raw data processing and normalization

In order to quantify the abundance of transcripts from 1,719 cells, salmon v0.17 ((Patro et al. 2017) was used. After indexing the human transcriptome (GRCh38.p13) in quasi-mapping-based mode, we quantified the transcripts with salmon, using --seqBias and --gcBias flags. We combined the transcript level abundances to the corresponding gene level counts, which were aggregated into a gene count matrix. Then, we filtered the cells based on the following criteria: we eliminated cells with less than 2,000 detected genes and overall mapping rate smaller than 55%. Cells with relatively high mapping rate to mitochondrial genes (>0.02) and ERCC spike-ins (>0.2) were also removed. After this step, we obtained 1,195 good quality cells. The data were normalized using ‘quickcluster’ and ‘normalize’ functions from scran package in R (Lun et al. 2016). This was followed by pseudocount addition of 1 and natural-log transformation of the count matrix.

### Clustering and cell type identi ication

To identify clusters of cells, we applied a graph-based algorithm. First, we selected the top 4,000 highly variable genes (HVGs) using the ‘high_variable_genes’ function from scanpy v1.4.4 69. We constructed the cell-cell distance matrix as 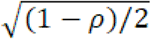, where *ρ* is the Spearman’s correlation coefficient between cells. Next, k-nearest neighbour graph was built with the first 30 principal components (PCs) and k=50. This was accomplished by the ‘neighbors’ function in scanpy, which computes the connectivity between cells based on UMAP 70. We chose the Leiden algorithm for community detection from the resulting graph, as it has been shown as a superior alternative to Louvain. We applied a resolution parameter of 0.75 to detect clusters in the data. The same resolution was used for sub-clustering the Endoderm and the Hemogenic Endothelial Progenitors clusters with top 2,000 HVGs in both. However, in this case the knn graph was built with the first 10 PCs and k=20. To visualize the resulting clusters in two dimensions, we computed a UMAP representation with default parameters on scanpy (‘tl.umap’ function).

We identified marker genes for a cluster with the Wilcoxon rank-sum test in scanpy (‘rank_genes_groups’ function) by comparing the gene expression levels within a given cluster with the rest of the cells. The genes were ranked according to FDR, after p-values were corrected with the Benjamini-Hochberg method. We visualized the expression values of marker genes on a heatmap, after scaling the log-normalized counts between 0 and 1 by using ‘standard_scale=var’ option in scanpy heatmap plotting function sc.pl.heatmap.

### Trajectory analysis using diffusion pseudotime and RNA-velocity

For the whole embryo diffusion map, we built the k-nearest neighbor graph as described above (with k=50 and using the first 30 PCs) to find the connectivity kernel width. We then used the ‘diffmap’ function to build the diffusion map.

To estimate the trajectory of epiblast differentiation, we took 2,000 HVGs from epiblast, primitive streak (PS), ectoderm and nascent mesoderm clusters combined. Finally, diffusion components were identified with k=15 and the first 15 PCs.

To illustrate the direction of differentiation in epiblast cells, we embedded the RNA velocities (La Manno et al. 2018) of single cells on the above diffusion map. For this task, we aligned reads from each cell using STAR v2.7 (Dobin et al. 2013) to the human reference genome (GRCh38.p13), which was obtained from ENSEMBL. The aligned bam files were processed with velocyto v0.17.17 (La Manno et al. 2018) with the default ‘run-smartseq2’ mode, to create a count matrix made of spliced and unspliced read counts.

After filtering genes with less than 10 spliced and un-spliced counts from this matrix, we calculated the moments for velocity estimation, by utilizing a built-in function from scVelo python module v01.20 (Bergen et al. 2019). Subsequently, we inferred the splicing kinetic dynamics of the genes by applying ‘recover_dynamics’ function. The velocity of each gene was estimated by solving splicing kinetics in the ‘dynamical’ mode with the ‘velocity’ function. Finally, we embedded the resulting velocities on the diffusion space calculated above by means of the ‘velocity_embedding’ function from the scVelo module. The diffusion map and the RNA velocities for the mesoderm specification analysis were computed in the same way.

We defined a diffusion pseudotime (dpt) coordinate on the diffusion map of epiblast differentiation, in order to visualize gene expression trends. First, we fixed the cell with the highest value of the first diffusion component (DC1) as root, so that the middle point of pseudotime would fall roughly into epiblast. We fitted the expression levels of the genes as a function of the pseudotime with a generalized additive model, by utilizing the ‘gam’ package in R (v1.16.1). For visualisation purpose, we transformed the pseudotime values as (1-dpt), so that ectoderm cells would fall onto the left side, and PS and nascent mesoderm on the right side of the pseudotime plot (Fig. 4a). Both fitted and unfitted values of the genes were scaled by dividing each by its maximum value of expression.

### Human-Mouse Comparison

For this analysis, we considered published single-cell RNA-seq data from mouse embryos during mid-streak stage (E7.25) (Pijuan-Sala et al. 2019), but the results remain largely unaffected if we use data from E7.0 or E7.5. Epiblast, primitive streak and nascent mesoderm clusters were selected from the human and the mouse datasets for downstream analysis, and they were analyzed separately as detailed below.

After constructing diffusion maps as described above with default parameters, we defined pseudotime starting from the cell with lowest DC1 value in both cases (Fig. S5a). After fitting gene expressions along pseudotime with generalized additive models (see above), we calculated the p-values using the ANOVA non-parametric test from the ‘gam’ R package. We then obtained the FDR values (Benjamini-Hochberg method). Genes with FDR<0.1 were clustered according to their expression pattern. This was achieved by hierarchical clustering with Spearman’s correlation distance as described above (‘hclust’ function in R). For estimating the number of clusters, the dynamic hybrid cut method was used (‘cutreeDynamic’ function, in the package dynamicTreeCut, version 1.63, with ‘deepslit’= 0 and ‘minclustersize’= 50). In both human and mouse, we found three clusters of genes, two of which were characterized by a clear upward or downward average trend with an absolute log-fold change greater than 1.

For the human-mouse comparison, we converted mouse genes to human equivalents (one-to-one homologous genes only) with the biomaRt package (Durinck et al. 2009). We compared the trends of genes in human and mouse, and in particular we looked at genes coding for signalling molecules, as listed in the curated database of the CellPhoneDB package (Efremova et al. 2020). To visualize the trend of selected genes, we normalized the expression values by the maximum in both mouse and human. We set fitted values to zero for the genes that were expressed in less than 10 cells.

### Blood staging analysis

Using the same mouse dataset, we selected epiblast, primitive streak, endothelium, blood progenitors (1 and 2), and erythroid (1, 2 and 3) clusters across the 9 stages, from E6.5 until E8.5. We merged the two blood progenitor clusters as well as 3 erythroid clusters and we obtained 4 mouse blood-related clusters that were used in downstream analyses. To build a representative expression pattern for each cluster/stage, we calculated the median expression value of the genes per cluster/stage. Cells from human gastrula blood (Erythrocytes, Myeloid Progenitors, Endothelium, Blood Progenitors, EMPs), epiblast and primitive streak clusters were projected onto the corresponding mouse clusters using the “scmapCluster” function from the “scmap” package in R with 1,000 genes and similarity threshold parameter set to 0.1.

### Primordial Germ Cell (PGC) identi ication

To single out the PGCs, we ran the algorithm called RaceID (“RaceID” package v0.1.5) (Grün et al. 2015), which can identify rare cell types, on the cells in the primitive streak cluster. We used these parameter values: k=1, outlg=8 and probthr=0.005. This resulted in the identification of 9 sub-clusters of outlier cells. Among these, the PGCs were identified as the only cluster of outlier cells that had a median expression of previously known PGC marker genes (NANOS3, SOX17, DND1, LAMA4, DPPA5) above 0.

### Cell cycle prediction

We estimated the cell cycle phase of each cell by applying the “pairs” algorithm, described by Scialdone et al. 2015 (Scialdone et al. 2015). A python implementation of this algorithm, ‘pypairs’ v3.1.1 was used in this analysis (https://pypairs.readthedocs.io/en/latest/documentation.html). After determining marker pairs from a training dataset (Leng et al. 2015) with the ‘sandbag’ function, we applied the function ‘cyclone’ to assign a cell cycle phase to each cell.

### Indel analysis

Using our transcriptomic data, we estimated the sizes of genomic insertions and deletions (indels) in our data as well as in a dataset from human fetal liver cells (Segal et al. 2019). This dataset was also processed with SMART-seq2 protocol and paired-end sequencing, although read lengths (75bp) were smaller than in our data (125bp). Hence, to minimize confounding effects in the results, we trimmed the reads in our data before processing it for this analysis. We aligned the data to the reference genome (GRCh38.p13), using bwa-mem v0.6 (Li 2013) with default parameters. We then merged the aligned data from each single cell into one bam file and performed indel calling with a pipeline for insertion and deletion detection from RNA-seq data called ‘transIndel’ v0.1 ((Yang et al. 2018). We kept the parameters at default values, except the minimum deletion length to be detected, which was set to 1 (-L flag set to 1).

### Differential gene expression analysis between rostral and caudal mesoderm

We used the R packages DESeq2 v3.11 (Love et al. 2014) and Seurat v3.0 (Stuart et al. 2019) to identify the genes differentially expressed between rostral and caudal parts of the mesoderm cluster. After creating a Seurat object with the mesoderm cells, their anatomical and plate information, we converted it to DESeq2 object with “convertTo” function. We found differentially expressed genes between caudal and rostral parts of the mesoderm with “DESeqDataSet” and “DESeq” functions, while controlling for the plate effect. Finally, we kept the genes with FDR < 0.1.

### Human embryonic stem cells comparison

For this comparison, we considered previously published single-cell RNA-seq data from pre-implantation human embryos (Petropoulos et al. 2016) and from hESC (Messmer et al. 2019). In the pre-implantation embryo data, we removed cells from extra-embryonic tissues, from immunosurgery samples and with unannotated stage. Moreover, we only kept cells with a log10 total number of reads greater than 5.5. This resulted in 442 cells distributed between E3 and E7 stages.

In the hESC dataset, only cells in batch 1 (including both primed and naïve hESC) that passed the quality test performed in the original publication were taken.

These data from pre-implantation embryos and hESC were combined with the epiblast cells in our dataset, and count per million (CPM) normalization was performed. We carried out a Principal Component Analysis (PCA) on the combined embryonic data using the top 1,600 highly variable genes (HVGs), and then we projected the hESC data onto this PCA.

To compare changes in gene expression levels between the naïve and primed state in epiblast and in hESC, we took cells from E6 stage, given that they showed the highest correlation with primed hESC (see Supp. Figure 5b). Then, the log-fold changes of the previously identified HVGs (after removal of genes with less than mean log count of 1) were calculated between E16 vs E6 cells and primed vs naïve hESC, after adding a pseudocount of 0.1 to the mean expression values. The line in Figure 3b is obtained through a linear regression (LinearRegression function from sklearn python module).

### Human gastruloid comparison

Recently published data from human gastruloids were considered for the comparison (Moris et al. 2020). Specifically, we considered the clusters of genes that exhibited a differential pattern of expression along the anterior-posterior axis of the gastruloids. After filtering out the clusters with less than 20 genes, we performed a statistical test to check whether the genes in each cluster were enriched for markers of the cell types we identified in the human gastrula.

To this aim, first we calculated the intersections between each gastruloid gene cluster and the list of marker genes of the cell types in the human gastrula:

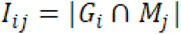

i.e. *I*_*ij*_, is the cardinality of the intersection between the i-th gastruloid gene cluster *G*_*i*_ and the list of marker genes of the j-th cell type in the human gastrula *M*_*j*_. We then computed the intersections with randomly shuffled lists of marker genes (keeping the number of markers per cluster constant and imposing that each gene could be listed at most once as marker of a given cluster):

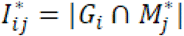

A null distribution for 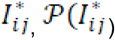,), was obtained by repeating the random shuffling 500 times. This allowed us to quantify the over-representation of the markers of a given cell type in each of the gastruloid gene cluster, by calculating:

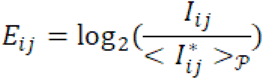

where 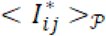 indicates the average value of 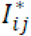 computed over the null distribution 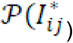. Additionally, a p-value *p_ij_* was estimated as:

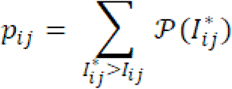

### Maintenance and differentiation of hESC

Human ESCs (H9/WA09 line; WiCell) were cultured on plates coated with 10 µg/ml vitronectin (Stem Cells Technologies) in 37°C and 5% CO2. Pluripotent hESCs were plated as single cells at 4.0-5.0×104 cells/cm2 using accutase (Gibco) and 10 µM Y27632 (Selleck), and maintained for two days in E6 media (Chen et al. 2011) supplemented with 2 ng/mL TGF-beta (bio-techne) and 25 ng/mL FGF2 (Dr. Marko Hyvönen, Cambridge University). These cells were sampled as “D0 PLU”. Then, the cells were cultured for one day in CDM/PVA media ((Johansson and Wiles 1995), 1 mg/ml polyvinyl alcohol (Sigma) instead of BSA) with 100 ng/ml Activin A (Dr. Marko Hyvönen, Cambridge University), 80 ng/ml FGF2, 10 ng/ml BMP4 (bio-techne), 10 µM LY294002 (Promega) and 3 mM CHIR99021 (Tocris), and sampled as “D1 ME” or “D1 ME+PD”. PD0325901 (Stem Cell Institute) was added at 1 µM. Bright field pictures were taken with Axiovert microscope (200M, Zeiss).

### Immunocytochemistry

Cells plated on vitronectin-coated round coverslips (Scientific Laboratory Supplies) were washed once with PBS, and fixed with 4% paraformaldehyde (Alfa Aesar) in PBS at RT for 10 min. Following another PBS wash, cells were incubated with 0.25% Triton in PBS at 4°C for 15-20 min, 0.5% BSA (Sigma) in PBS at room temperature for 30 min, primary antibodies at 4°C overnight and secondary antibodies at room temperature for one hour. Anti-Ecadherin antibody (3195, Cell Signaling Technology, 1:200), and anti-Rabbit IgG Alexa Fluoro 568 antibody (A10042, Invitrogen, 1:1000) together with 10 µg/ml Hoechst33258 (B2883) were diluted in 0.5% BSA in PBS and each staining was followed by three washes with 0.5% BSA in PBS. Coverslips were preserved on slide glasses (Corning) with ProLong Gold Antifade Mountant (Life Technologies) and nail polisher, and observed with Zeiss inverted confocal system (LSM 710, Zeiss).

### Quantitative RT-PCR for hESC samples

Total RNA was extracted from cells using the GenElute Mammalian Total RNA Miniprep Kit (Sigma-Aldrich) and the On-Column DNase I Digestion set (Sigma-Aldrich). Complementary DNA was synthesized from the RNA using random primers (Promega), dNTPs (Promega), RNAseOUT (Invitrogen) and SuperScript II (Invitrogen). Real-time PCR was performed with KAPA SYBR FAST qPCR Master Mix (Kapa Biosystems) on QuantStudio 12K Flex Real-Time PCR System machine (Thermo Fisher Scientific). Molecular grade water (Thermo Fisher Scientific) was used when necessary. Each gene expression level was normalized by the average expression level of PBGD and RPLP0. Primer sequences are shown in Supplementary Table 7.

## AUTHOR CONTRIBUTIONS

RT, SN and SS performed the experiments; EM, and AS performed the computational analyses; RT, EM, SN, LV, AS and SS analysed the results and wrote the manuscript.

## ACKNOWLEDGEMENTS

Human embryonic material was provided by the MRC/Wellcome Trust funded (grant # 099175/Z/12/Z and MR/ R006237/1) Human Developmental Biology Resource (www.hdbr.org). We thank Neil Ashley (Oxford MRC single cell facility) for help with sequencing, Marella De Bruijn, Bertie Gottgens, Liz Robertson, Tristan Rodriguez and Maria-Elena Torres-Padilla for helpful comments. This work was funded by: British Heart Foundation Immediate Postdoctoral Basic Science Research Fellowship no. FS/18/24/33424 to RT; JSPS Overseas Research Fellowship to SN; European Research Council advanced grant ERC: 741707 to LV; funding from the Helmholtz Association to AS; Wellcome Strategic Awards 105031/C/14/Z, 108438/Z/15/Z, and Senior Investigator Award 103788/Z/14/Z to SS.

---

**Supplementary Figure 1.**
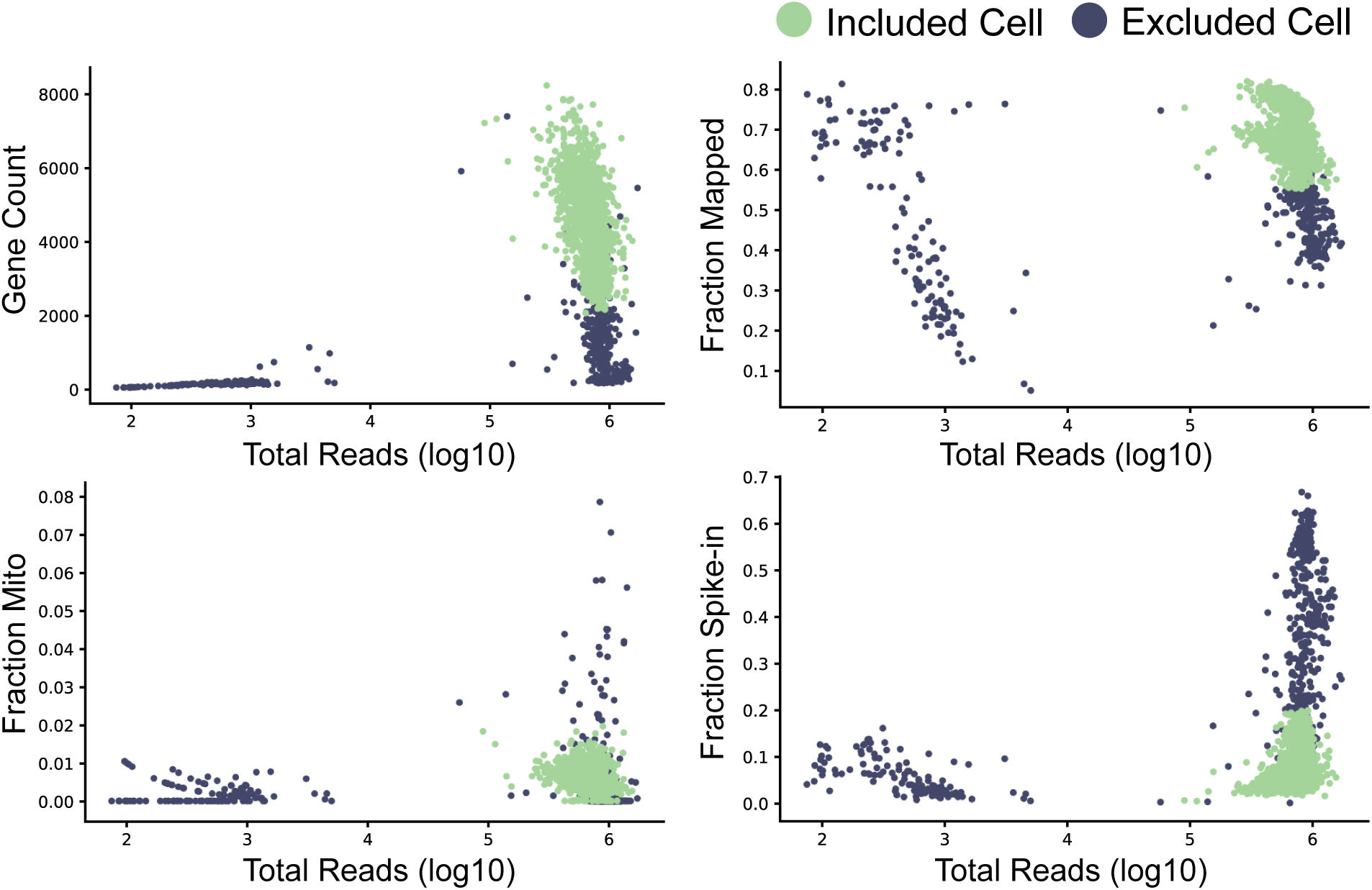
Quality control of scRNA-seq dataset. Metrics used to assess the quality of the scRNA-seq libraries. The scatter plots show the number of detected genes (top left), the fraction of reads mapped to the human genome (top right), the fraction of reads mapped to mitochondrial genes (bottom left) and the fraction of reads mapped to ERCC spike-ins (bottom right), all as a function of the total number of reads. Cells that passed quality control are marked by green circles, while black circles indicate cells that failed the quality control and were excluded from downstream analyses.

**Supplementary Figure 2.**
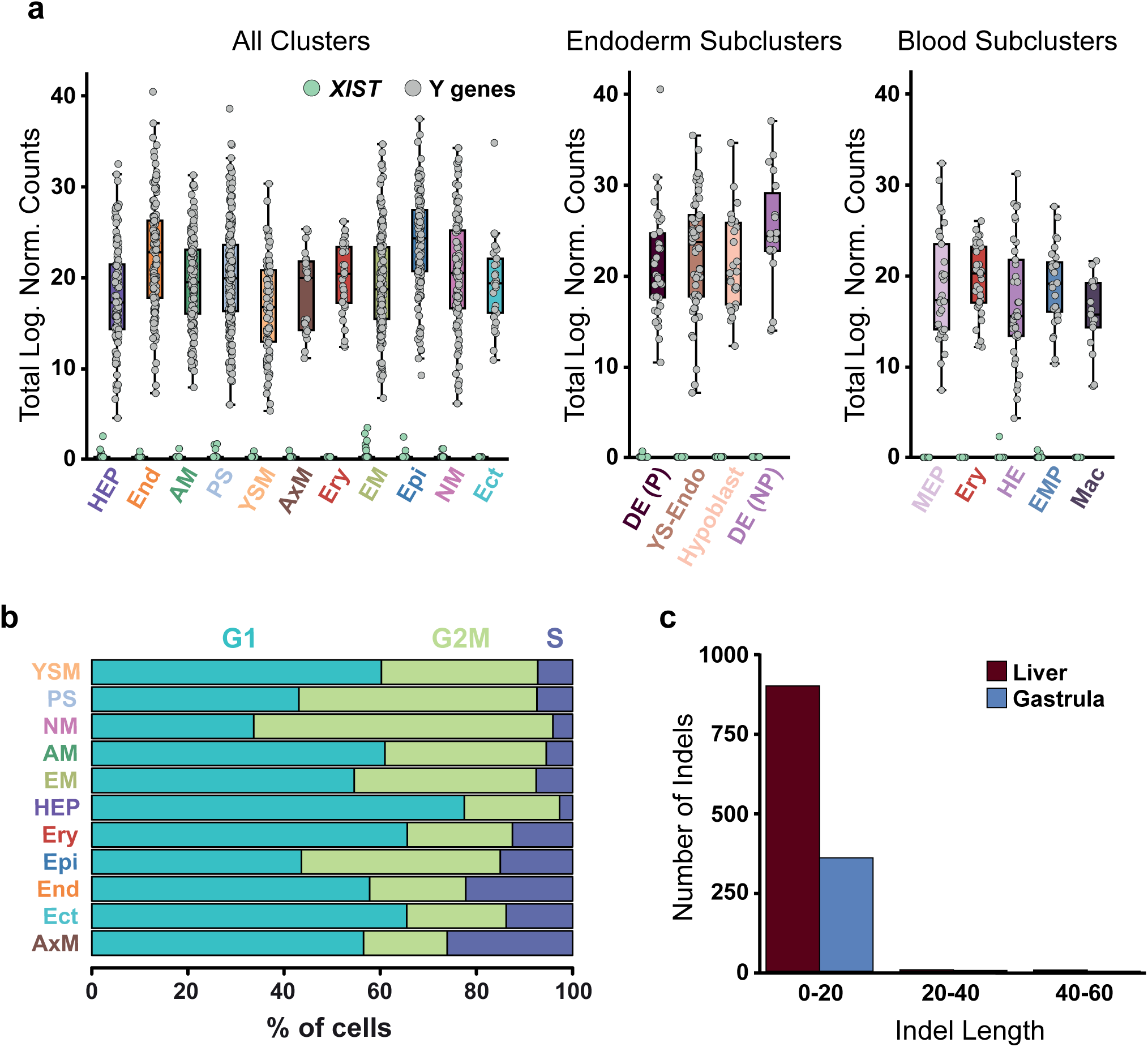
Maternal contamination and cell cycle analysis. a, The boxplots show the total log expression of normalized counts for XIST and Y-genes across all clusters (left), endodermal sub clusters (centre) and blood sub clusters (right). While XIST was mostly not detected, Y-chromosome genes had always non-zero counts; this suggests that there is no contamination from maternal tissues in any of the clusters. b, The stacked barplots indicate the percentages of cells from each cluster in the phase G1, S or G2M of the cell cycle, as predicted from their transcriptomic profiles. Clusters are Hemogenic Endothelial Progenitors (HEP), Endoderm (End), Advanced Mesoderm (AM), Primitive Streak (PS), Yolk Sac Mesoderm (YSM), Axial Mesoderm (AxM), Erythrocytes (Ery), Emergent Mesoderm (EM), Epiblast (Epi), Nascent Mesoderm (NM), Ectoderm (Ect). Endodermal sub clusters are Proliferative Definitive endoderm (DE(P)), Yolk Sac Endoderm (YS-Endo), Hypoblast and Non-Proliferative Definitive Endoderm (DE(NP)). Blood sub-clusters are Megakaryocyte-Erythroid progenitors (MEP), Erythrocytes (Ery), Hemogenic Endothelium (HE), Erythroid-Myeloid progenitors (EMPs) and Macrophages (Mac). c, Insertion-deletion length and size distribution of gastrula and fetal liver data. Y axis represents total number of indels on merged cells, while x axis represents indel length in base pairs.

**Supplementary Figure 3.**
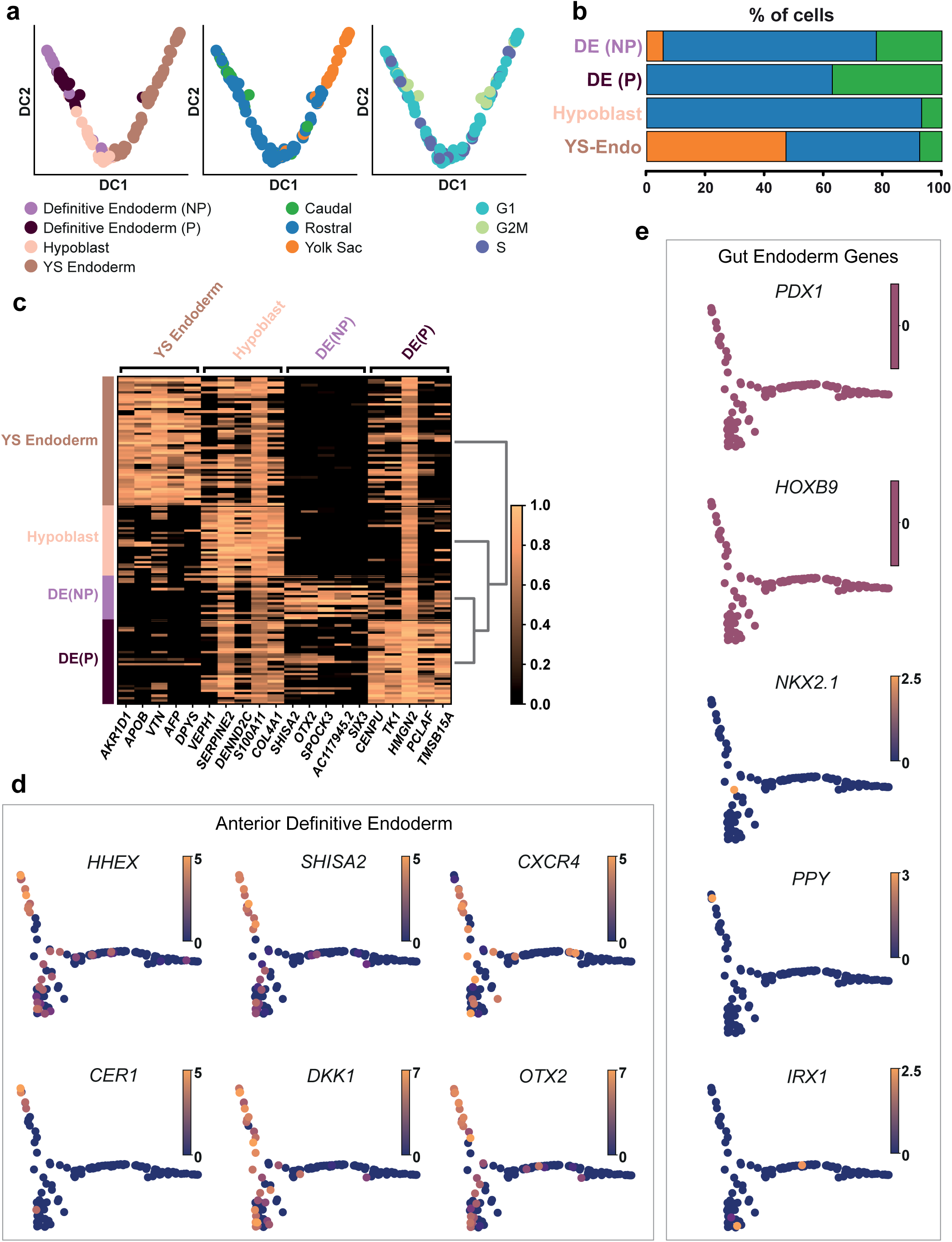
Endoderm subcluster identification. a, Diffusion map of cells from the Endoderm clusters. The first two diffusion components (DC1 and DC2) are plotted and cells are coloured by the sub clusters (left panel), the anatomical region of origin (central panel) or the predicted cell-cycle phase (right panel). Yolk Sac, YS;Non-Proliferative, NP; Proliferative, P. b, Percentage of cells dissected from the Caudal, Rostral or Yolk Sac portion of the embryo in the four endodermal sub-clusters. c, Heatmap showing the standardized log expression levels of marker genes of the four endodermal sub-clusters. d, Diffusion map of cells from the Endoderm clusters (same as in Figure 2) with cells coloured by the normalized log expression levels of Anterior Definitive Endoderm markers. These genes are more highly expressed in the Non-Proliferative Definitive Endoderm cluster. e, Diffusion map of cells from the Endoderm clusters (same as in Figure 2) with cells coloured by the normalized log expression levels of Gut Endoderm markers, showing limited expression.

**Supplementary Figure 4.**
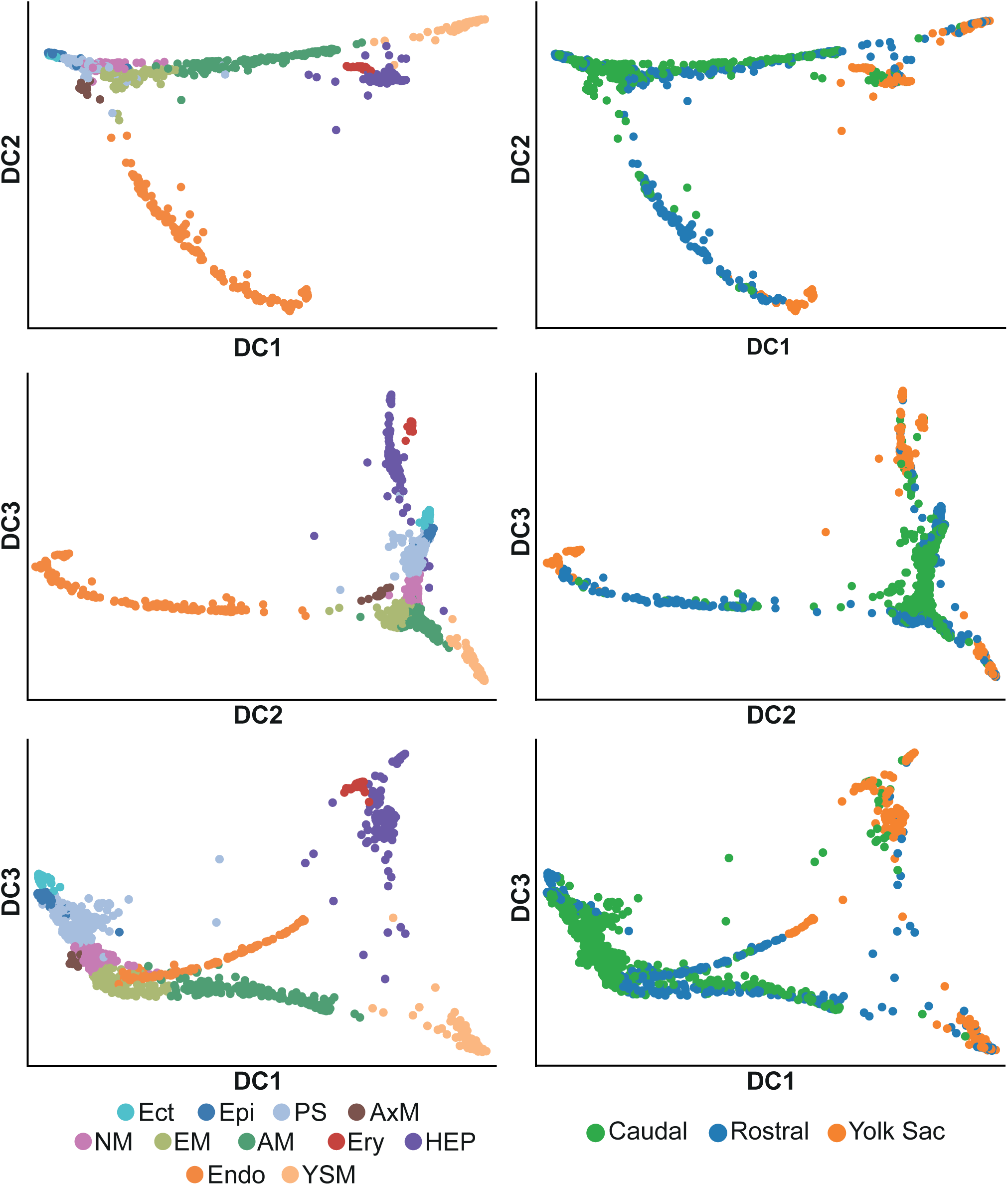
Diffusion maps for entire dataset. Diffusion map of cells from all 11 clusters, with the second (DC2) and the third (DC3) (top panels) or the first (DC1) and the third diffusion component shown (bottom panels). In the left panels cells are coloured by the clusters they belong to, while in the right panel the colours indicate the region each cell was dissected from. Ectoderm (Ect), Epiblast (Epi), Primitive Streak (PS), Axial Mesoderm (AxM), Nascent Mesoderm (NM), Emergent Mesoderm (EM), Advanced Mesoderm (AM), Erythrocytes (Ery), Hemogenic Endothelial Progenitors (HEP), Endoderm (Endo), Yolk Sac Mesoderm (YSM).

**Supplementary Figure 5.**
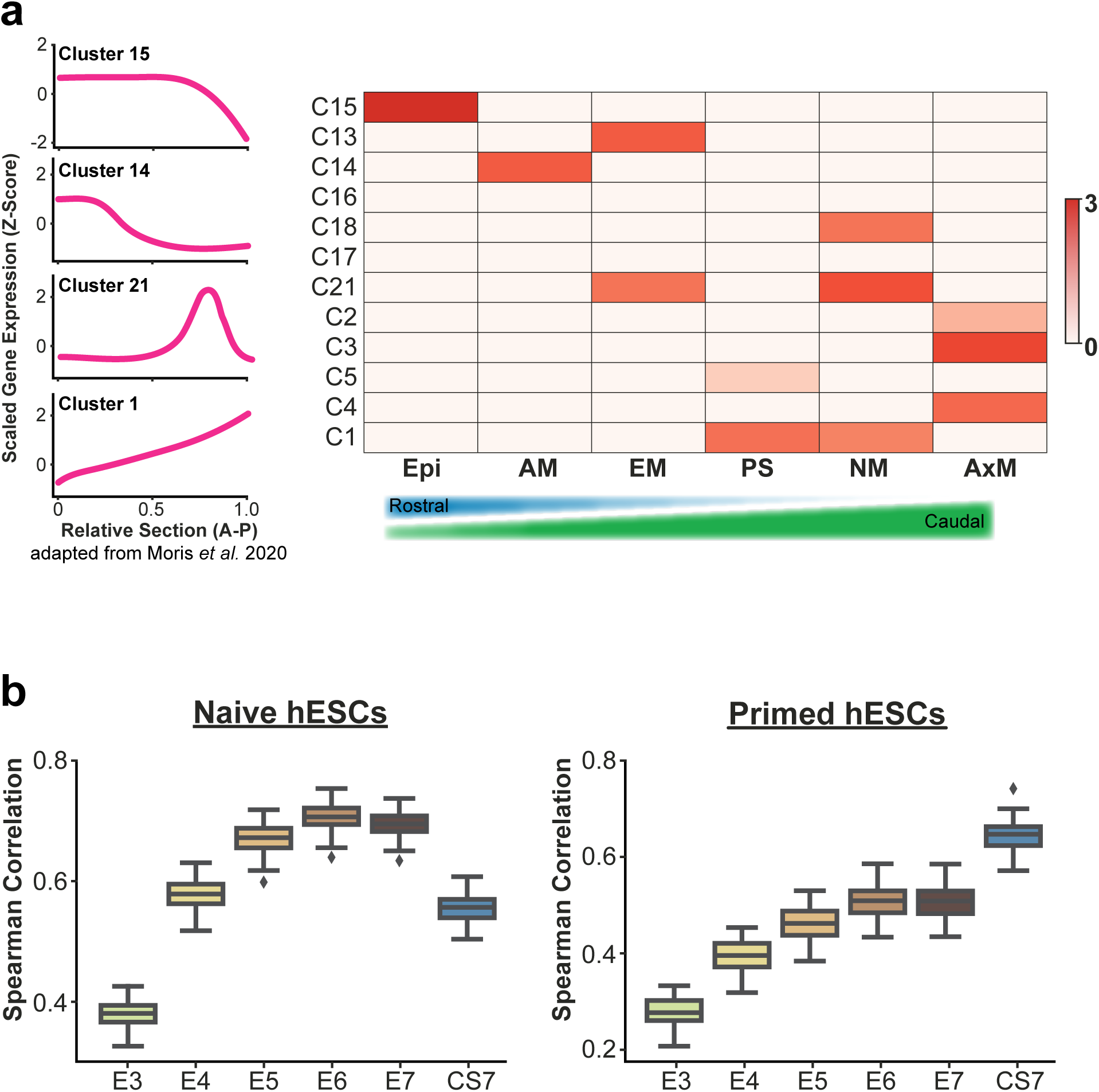
In Vitro vs In Vivo comparisons. a, In each cluster of genes with similar patterns along the anterior to posterior axis of human organoids (Moris et al. 2020) (indicated in rows), we calculated the log2-fold enrichment of markers for the human gastrula cell types (in columns; see Methods). In this heatmap, the enrichments that are statistically significant (p-value < 0.01; see Methods) are shown. The gastruloid gene clusters are sorted based on the approximate position of their peak of expression: from the most anterior at the top, to the most posterior at the bottom. Similarly, the human gastrula cell type are sorted based on the relative fraction of cells from the rostral and caudal region, as indicated by the triangles at the bottom: the clusters with more rostral cells are on the left, whereas those with more cells coming from the caudal region are on the right. b, Spearman’s correlation coefficients between the transcriptome of naïve (left) and primed (right) hESC and cells from the embryonic lineage of pre-implantation human embryos (from embryonic day (E) 3 to 7) and from the epiblast cells of a CS7 human gastrula. In particular, the boxplots show the correlation coefficients between each hESC and the average gene expression profiles of the cells from embryos at the different stages. Only highly variable genes computed on the embryonic data were used (see Methods).

**Supplementary Figure 6.**
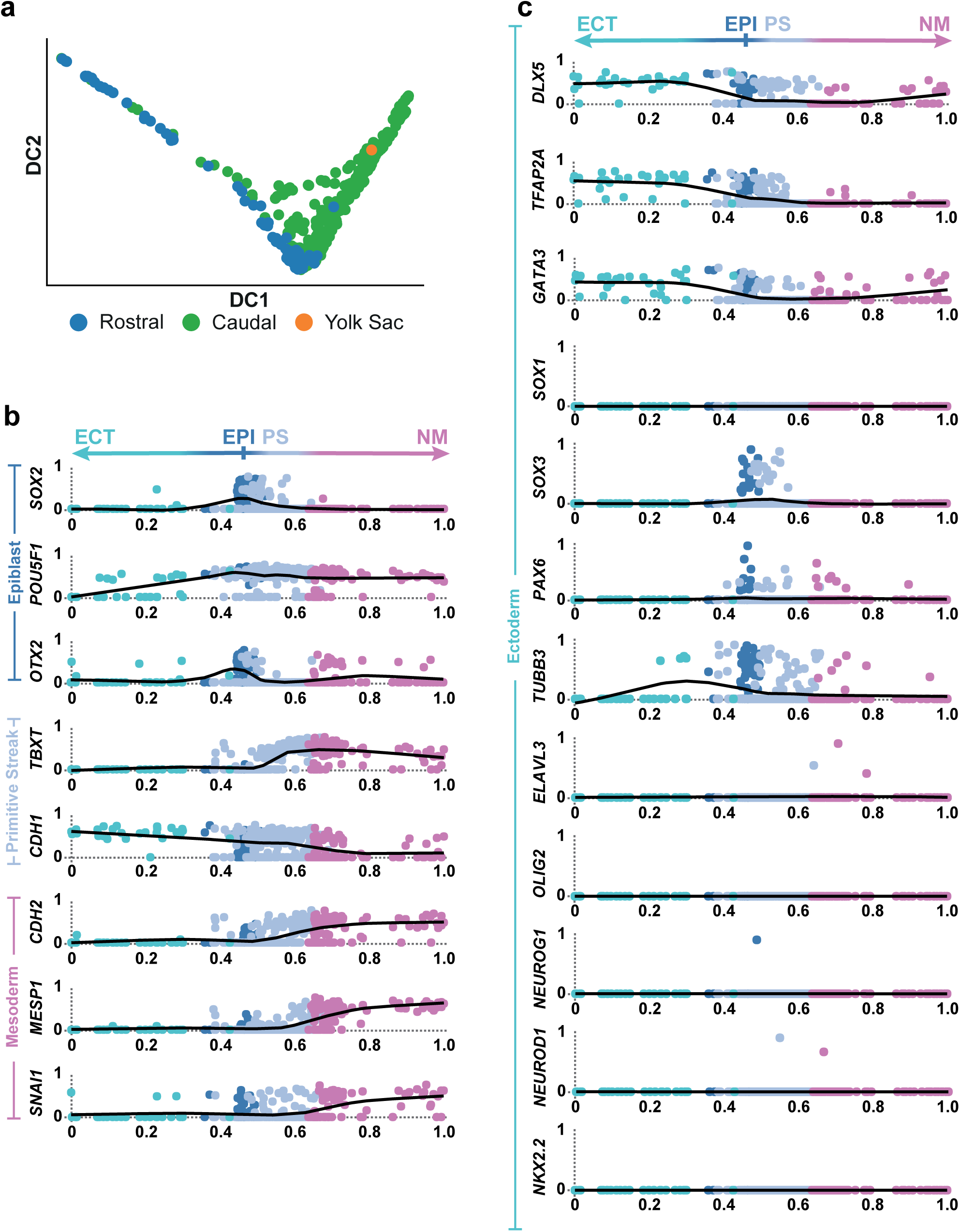
Differentiation of epiblast. a, Diffusion map of cells from the Epiblast, Ectoderm, Primitive Streak and Nascent Mesoderm (same as in Figure 4a). The first two diffusion components are plotted (DC1 and DC2) and cells are colored by the anatomical region they were isolated from. b, c Standardized gene expression changes along a pseudotime coordinate (see Figure 4a) running from 0 to 1 and spanning the Ectoderm (ECT), the Epiblast (EPI), the Primitive Streak (PS) and the Nascent Mesoderm (NM), as depicted by the arrow on top. The selected genes highlight Primitive Streak and mesoderm formation (panel b) as well as ectoderm differentiation (panel c).

**Supplementary Figure 7.**
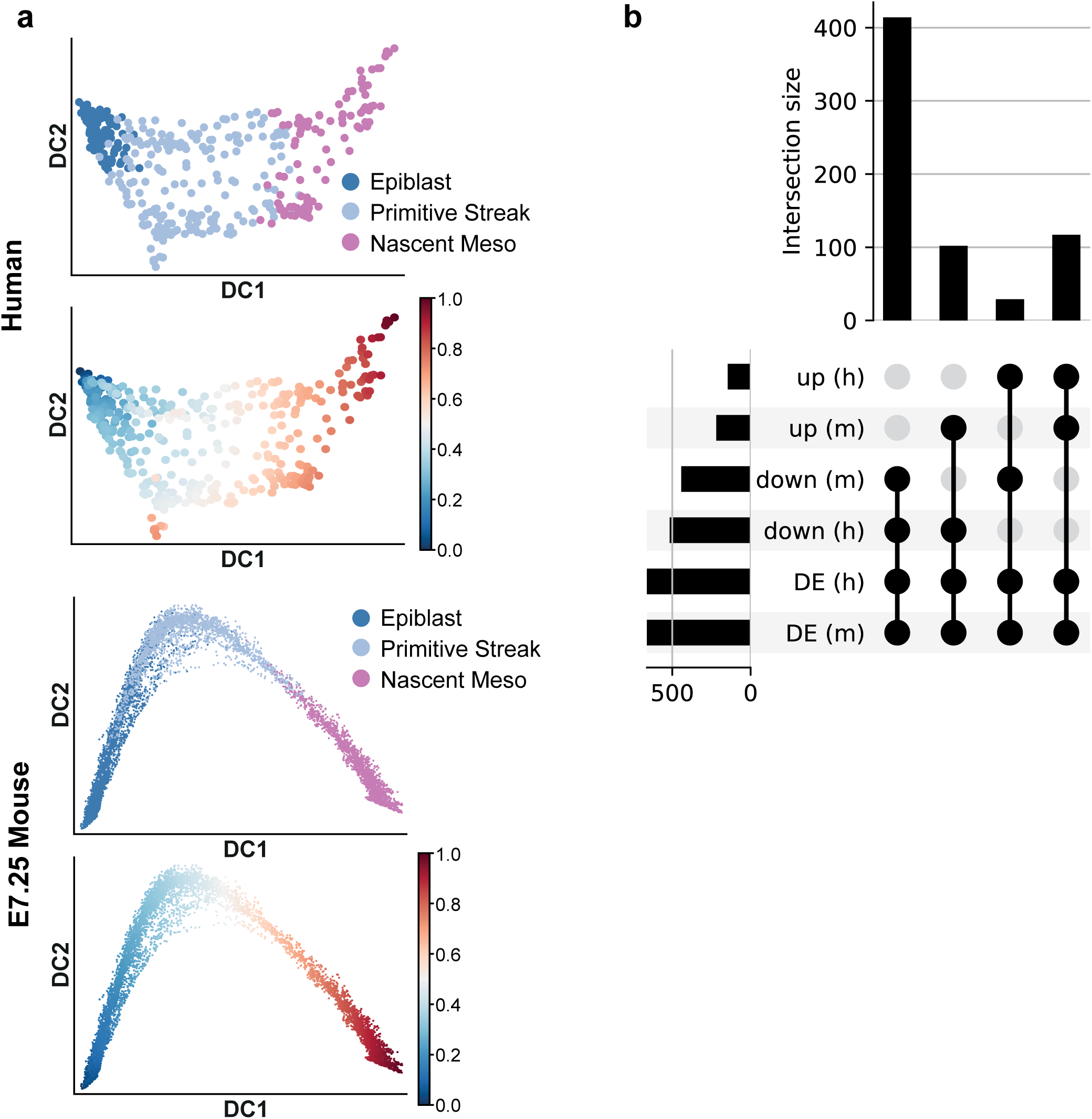
Mesoderm formation in human and mouse. a, Diffusion map with cells from the human (top two plots) or mouse (bottom two plots) Epiblast, Primitive Streak and Nascent Mesoderm clusters. Cells are colored based on their cluster of origin or on their diffusion pseudotime coordinate. b, Upset plot for the number of differentially expressed (DE) genes as a function of the diffusion pseudotime (dpt) shown in panel a in mouse (m) or human (h). Genes are split according to their increasing (up) or decreasing (down) trend as a function of dpt.

**Supplementary Figure 8.**
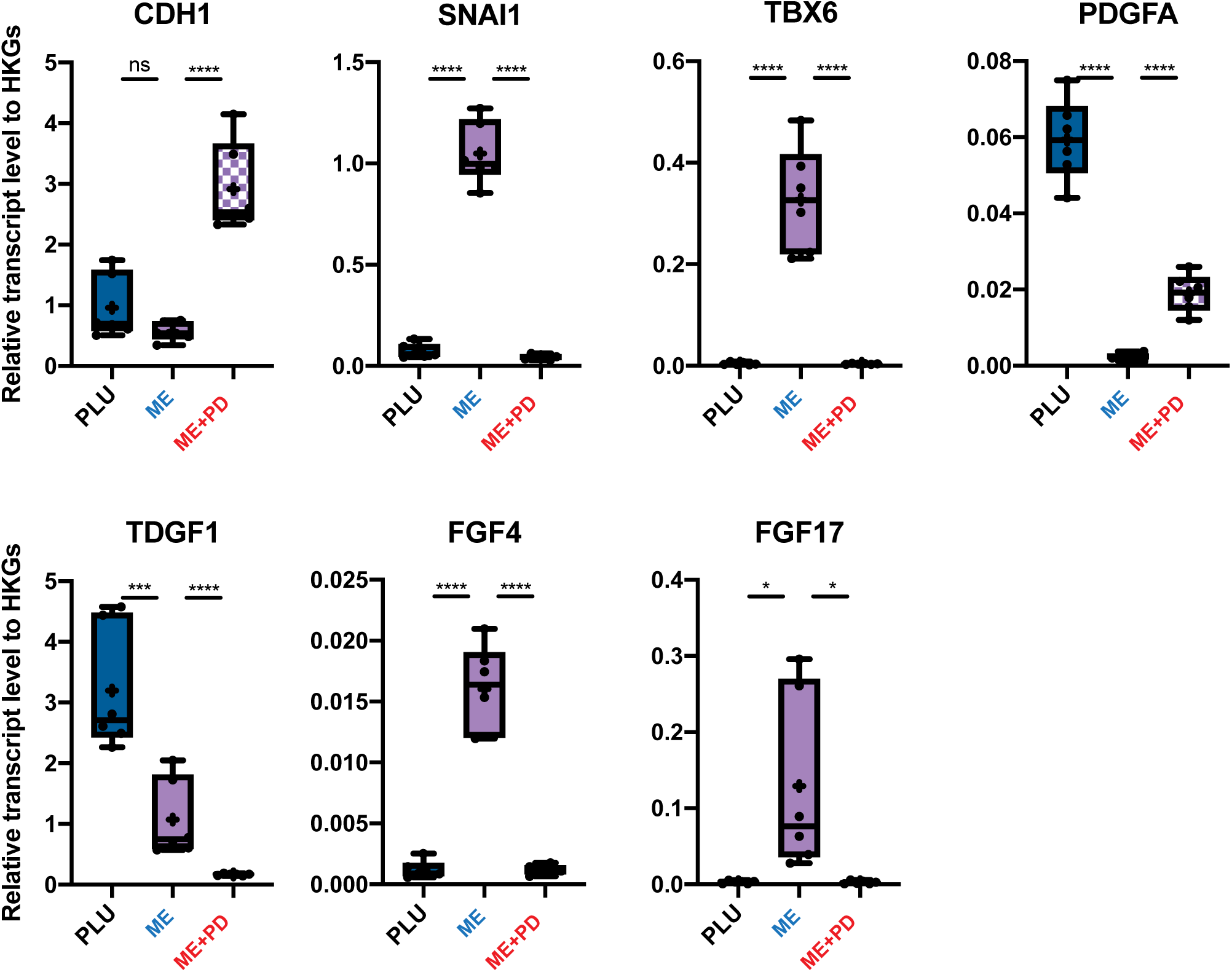
Characterization of EMT during hESC mesoderm formation. Quantification of transcript levels for selected genes across the three conditions PLU, ME, ME+PD. (n = 6 from three different experiments. ns = p-value ≥ 0.05; *** = p-value < 0.001; **** = p-value < 0.0001 (Ordinary one-way ANOVA after Shapiro-Wilk normality test).

**Supplementary Figure 9.**
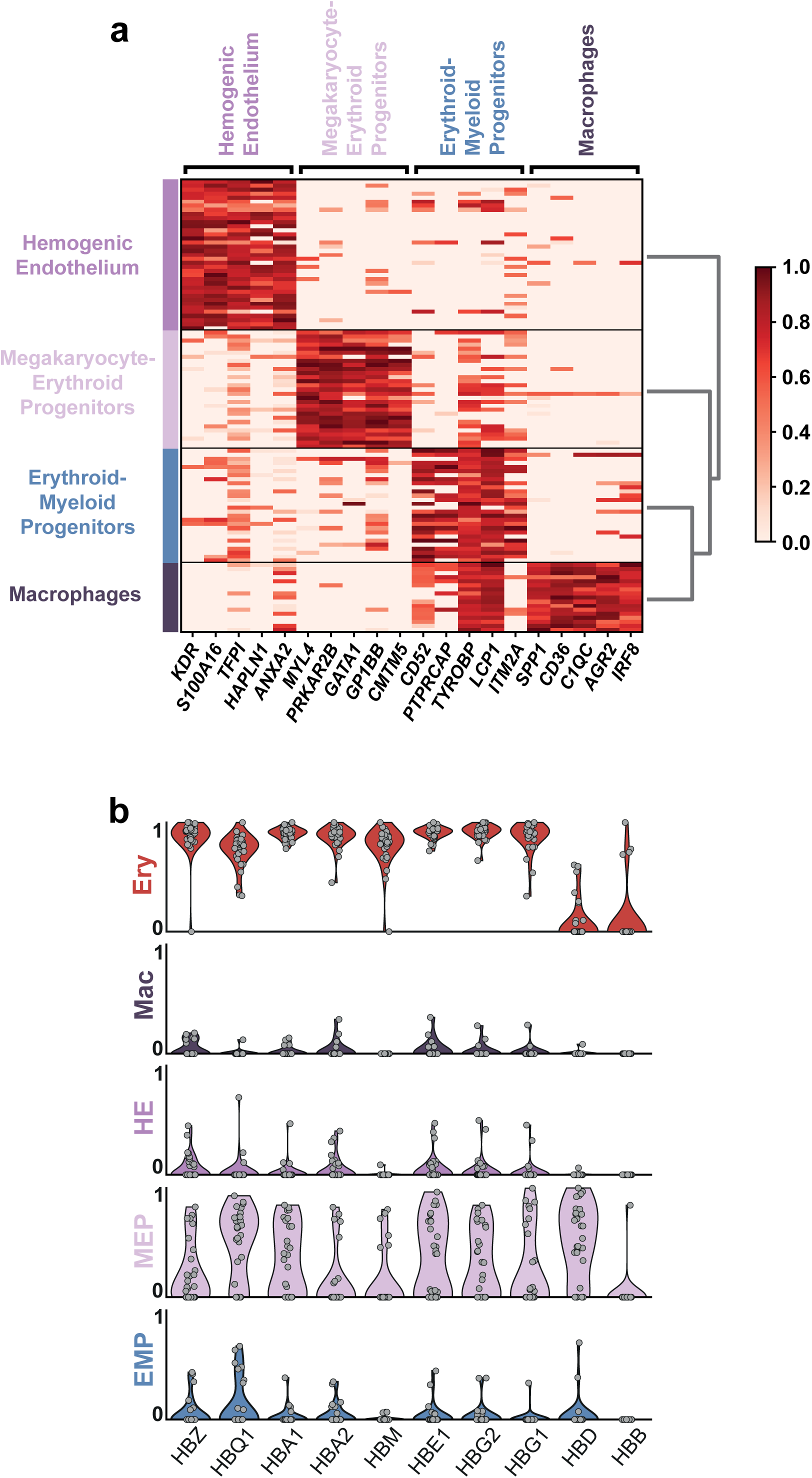
Hemogenic Endothelial Progenitors subclusters. a, Heatmap showing the standardized log expression levels of the top 5 marker genes of the four Hemogenic Endothelial Progenitors sub-clusters. b, Violin plots showing the normalized expression of Hemoglobin genes in the five blood related clusters: Erythrocytes (Ery), Macrophages (Mac), Hemogenic Endothelium (HE), Megakaryocyte-Erythroid progenitors (MEP) and Erythroid-Myeloid progenitors (EMPs). Each grey dot represents a single cell.

**Supplementary Table 1.**
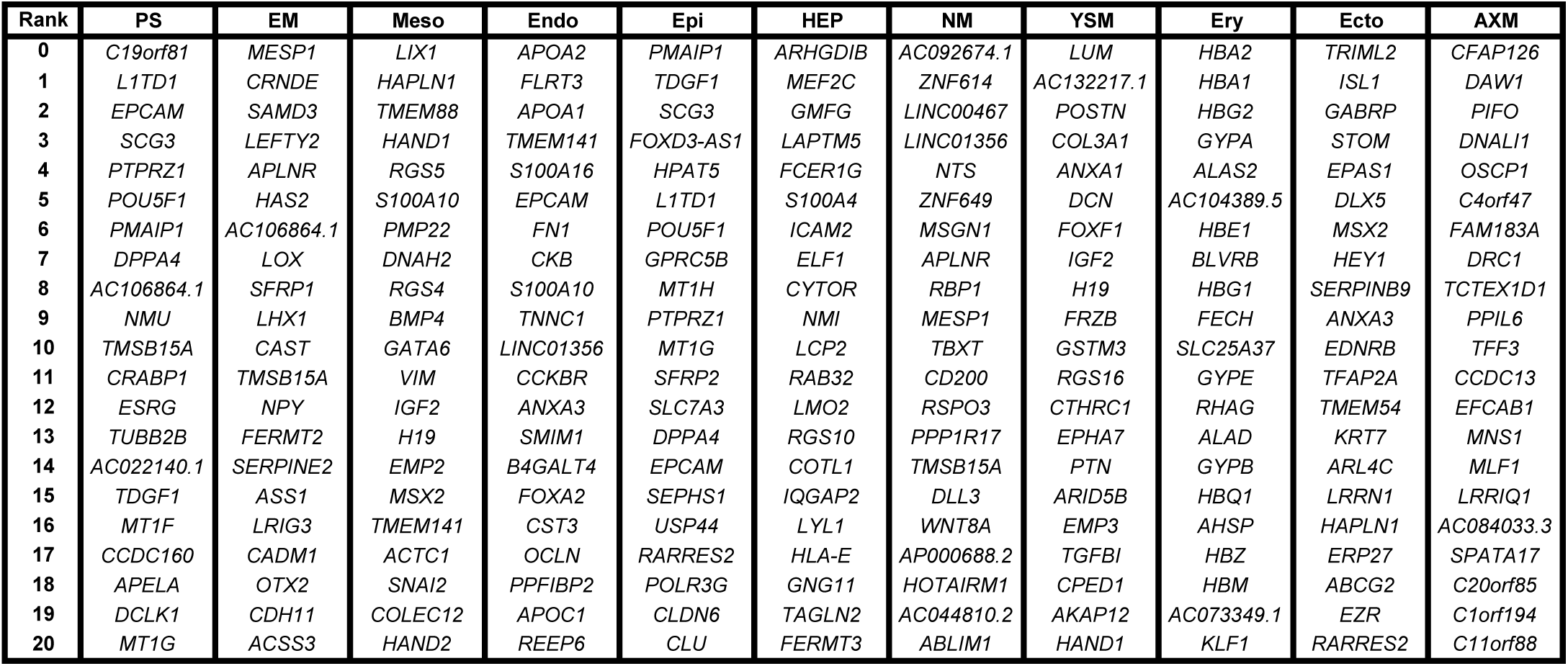
Human Gastrula Cluster Genes.

**Supplementary Table 2.** Cell Orgin per Cluster.

| Cluster | Origin | Percentage |
| --- | --- | --- |
| Axial Mesoderm | Caudal | 100.0 |
| Ectoderm | Caudal | 20.7 |
|  | Rostral | 79.3 |
| Endoderm | Caudal | 17.0 |
|  | Rostral | 63.7 |
|  | Yolk Sac | 19.3 |
| Epiblast | Caudal | 54.9 |
|  | Rostral | 45.1 |
| Erythrocytes | Caudal | 18.8 |
|  | Yolk Sac | 81.3 |
| Hemogenic<br>Endothelial<br>Progenitors | Caudal | 15.3 |
|  | Rostral | 12.6 |
|  | Yolk Sac | 72.1 |
| Immature Mesoderm | Caudal | 69.7 |
|  | Rostral | 30.3 |
| Mature Mesoderm | Caudal | 57.9 |
|  | Rostral | 42.1 |
| Nascent Mesoderm | Caudal | 99.0 |
|  | Yolk Sac | 1.0 |
| Primitive Streak | Caudal | 96.0 |
|  | Rostral | 4.0 |
| YS Mesoderm | Caudal | 2.4 |
|  | Rostral | 28.9 |
|  | Yolk Sac | 68.7 |

**Supplementary Table 3.** Top 20 Primordial Germ Cell DEGs.

| **Rank** | **PGC** | **FDR** | **Primitive Streak** | **FDR** |
| --- | --- | --- | --- | --- |
| **0** | *NANOG* | 9.31E-03 | *HMGN2* | 9.31E-03 |
| **1** | *FAM162B* | 9.31E-03 | *MDK* | 9.31E-03 |
| **2** | *TCL1A* | 9.31E-03 | *SRP9* | 9.31E-03 |
| **3** | *S100A10* | 9.31E-03 | *GNAS* | 9.31E-03 |
| **4** | *LAMA4* | 9.31E-03 | *PPIA* | 9.31E-03 |
| **5** | *DPPA5* | 9.31E-03 | *BEX3* | 9.55E-03 |
| **6** | *IFI16* | 9.31E-03 | *TSTD1* | 1.11E-02 |
| **7** | *NANOS3* | 9.31E-03 | *PTMA* | 1.16E-02 |
| **8** | *PCSK1N* | 9.31E-03 | *HMGB1* | 1.16E-02 |
| **9** | *RASD1* | 9.31E-03 | *HSBP1* | 1.16E-02 |
| **10** | *SDCBP* | 9.55E-03 | *DEK* | 1.16E-02 |
| **11** | *SAT1* | 1.16E-02 | *FTL* | 1.16E-02 |
| **12** | *FAM122C* | 1.16E-02 | *GSTO1* | 1.16E-02 |
| **13** | *PDPN* | 1.16E-02 | *PCLAF* | 1.16E-02 |
| **14** | *ASRGL1* | 1.16E-02 | *LDHB* | 1.16E-02 |
| **15** | *HERC5* | 1.42E-02 | *ARRDC3* | 1.16E-02 |
| **16** | *MKRN1* | 1.56E-02 | *RANBP1* | 1.16E-02 |
| **17** | *CDK2AP1* | 1.86E-02 | *UBE2V2* | 1.16E-02 |
| **18** | *SOX15* | 2.18E-02 | *SINHCAF* | 1.16E-02 |
| **19** | *IFITM1* | 2.18E-02 | *PRDX2* | 1.16E-02 |
| **20** | *RMND1* | 2.28E-02 | *TMSB15A* | 1.16E-02 |

**Supplementary Figure 4.** Top20 DEG during human Epiblast to Nascent mesoderm differentiation.

| **Rank** | **Upregulated** | **FDR** | **Downregulated** | **FDR** |
| --- | --- | --- | --- | --- |
| 1 | *APLNR* | 4.63E-98 | *TDGF1* | 2.46E-68 |
| 2 | *AC092674.1* | 7.16E-96 | *MT1G* | 5.17E-64 |
| 3 | *LINC01356* | 9.97E-67 | *ADM* | 1.40E-60 |
| 4 | *MESP1* | 3.08E-55 | *MT2A* | 2.30E-59 |
| 5 | *RSPO3* | 4.57E-54 | *MT1H* | 2.01E-58 |
| 6 | *MSGN1* | 5.21E-54 | *EPCAM* | 3.71E-50 |
| 7 | *RBP1* | 7.16E-45 | *PODXL* | 1.33E-49 |
| 8 | *CDH2* | 1.39E-42 | *HPAT5* | 2.83E-49 |
| 9 | *CDH11* | 1.75E-41 | *PMAIP1* | 8.24E-46 |
| 10 | *TBXT* | 1.75E-41 | *MT1E* | 2.29E-44 |
| 11 | *LINC00467* | 3.55E-41 | *MAL2* | 4.35E-44 |
| 12 | *KCNK17* | 6.20E-37 | *MT1F* | 9.92E-44 |
| 13 | *WNT8A* | 6.48E-37 | *PHLDA1* | 2.68E-42 |
| 14 | *DLL3* | 4.12E-36 | *CLU* | 4.41E-42 |
| 15 | *ENPP2* | 7.93E-35 | *GPRC5B* | 3.26E-41 |
| 16 | *PPP1R17* | 1.19E-33 | *VAT1L* | 6.09E-41 |
| 17 | *HVCN1* | 1.84E-28 | *FOXD3-AS1* | 1.43E-40 |
| 18 | *PDGFRA* | 6.26E-28 | *APELA* | 5.23E-38 |
| 19 | *AC073174.2* | 1.38E-26 | *SLC7A3* | 7.71E-36 |
| 20 | *PPIAP46* | 2.88E-26 | *BIK* | 2.28E-35 |

**Supplementary Figure 5.** Top20 DEG during mouse Epiblast to Nascent mesoderm differentiation.

| **Rank** | **<u>Upregulated</u>** | **<u>FDR</u>** | **<u>Downregulated</u>** | **<u>FDR</u>** |
| --- | --- | --- | --- | --- |
| 1 | *APLNR* | 4.63E-98 | *EPCAM* | 3.71E-50 |
| 2 | *TDGF1* | 2.46E-68 | *SLC7A3* | 7.71E-36 |
| 3 | *MESP1* | 3.08E-55 | *ENPP2* | 7.93E-35 |
| 4 | *RSPO3* | 4.57E-54 | *PPP1R1A* | 1.32E-34 |
| 5 | *RBP1* | 7.16E-45 | *SOX2* | 7.15E-32 |
| 6 | *CDH2* | 1.39E-42 | *CLDN6* | 1.31E-29 |
| 7 | *PHLDA1* | 2.68E-42 | *ARL4A* | 7.35E-29 |
| 8 | *TBXT* | 1.75E-41 | *HVCN1* | 1.84E-28 |
| 9 | *CDH11* | 1.75E-41 | *USP44* | 2.75E-28 |
| 10 | *DLL3* | 4.12E-36 | *SLC29A1* | 5.47E-27 |
| 11 | *PDGFRA* | 6.26E-28 | *UCHL1* | 1.66E-26 |
| 12 | *SOCS2* | 3.32E-27 | *RAB25* | 2.91E-25 |
| 13 | *CNMD* | 9.02E-27 | *KIF1A* | 1.11E-22 |
| 14 | *EGLN3* | 8.93E-26 | *SALL2* | 7.01E-22 |
| 15 | *TBX6* | 5.93E-24 | *BCAT2* | 9.06E-22 |
| 16 | *HOXB2* | 1.14E-23 | *PHC1* | 5.11E-21 |
| 17 | *FGF3* | 2.68E-22 | *DNMT3B* | 8.43E-21 |
| 18 | *MEIS2* | 1.72E-21 | *CLDN3* | 1.65E-20 |
| 19 | *WNT5A* | 5.50E-21 | *ID3* | 1.71E-20 |
| 20 | *ID1* | 9.22E-21 | *ESRP1* | 5.61E-20 |

**Supplementary Table 6.** Top 20 HEP Subcluster Genes.

| Rank | EMP | FDR | HE | FDR | Myeloid Progenitors | FDR | Blood Progenitors | FDR |
| --- | --- | --- | --- | --- | --- | --- | --- | --- |
| 0 | CD52 | 1.16E-04 | KDR | 2.58E-13 | SPP1 | 4.53E-07 | MYL4 | 1.72E-11 |
| 1 | PTPRCAP | 3.19E-04 | S100A16 | 8.11E-12 | CD36 | 4.53E-07 | PRKAR2B | 1.72E-11 |
| 2 | TYROBP | 7.67E-04 | TFPI | 1.13E-11 | C1QC | 4.53E-07 | GATA1 | 9.57E-11 |
| 3 | LCP1 | 9.71E-04 | HAPLN1 | 1.50E-11 | AGR2 | 5.52E-07 | GP1BB | 1.32E-10 |
| 4 | ITM2A | 1.39E-03 | ANXA2 | 2.53E-11 | IRF8 | 6.09E-07 | CMTM5 | 1.32E-10 |
| 5 | CYTL1 | 2.87E-03 | EFNA1 | 6.80E-11 | CCL3 | 6.98E-07 | ITGA2B | 3.73E-10 |
| 6 | CSF3R | 2.93E-03 | ECSCR | 2.82E-10 | TIMD4 | 7.01E-07 | GP1BA | 1.48E-09 |
| 7 | ATP8B4 | 4.21E-03 | PLK2 | 7.22E-10 | RASSF4 | 7.56E-07 | GP9 | 1.48E-09 |
| 8 | SPINK2 | 4.21E-03 | PHLDA1 | 8.54E-10 | SAMHD1 | 9.69E-07 | LY6G6F | 1.48E-09 |
| 9 | GUCY1A1 | 4.21E-03 | S100A10 | 8.54E-10 | A2M | 1.19E-06 | CCND3 | 1.70E-09 |
| 10 | AIF1 | 5.15E-03 | ID3 | 1.65E-09 | ABCG2 | 1.90E-06 | RHAG | 2.07E-09 |
| 11 | CD200 | 7.38E-03 | FSTL1 | 3.96E-09 | FTL | 2.29E-06 | TUBB1 | 2.31E-09 |
| 12 | ZFP36L2 | 8.26E-03 | VAMP5 | 8.36E-09 | C1QA | 2.29E-06 | CD55 | 3.69E-09 |
| 13 | JAML | 9.56E-03 | MDK | 1.28E-08 | CYBB | 2.43E-06 | SMIM1 | 4.13E-09 |
| 14 | S100A4 | 1.07E-02 | SPARC | 1.68E-08 | MTUS1 | 2.43E-06 | C2orf88 | 4.13E-09 |
| 15 | ARHGAP25 | 1.18E-02 | RAMP2 | 1.78E-08 | CD14 | 2.45E-06 | LY6G6D | 5.00E-09 |
| 16 | CD44 | 1.35E-02 | CRIP2 | 4.68E-08 | LSP1 | 2.61E-06 | NFE2 | 5.00E-09 |
| 17 | PKIB | 1.35E-02 | ARHGAP29 | 5.76E-08 | MILR1 | 2.87E-06 | THBS1 | 5.00E-09 |
| 18 | SMIM24 | 1.36E-02 | CALCRL | 5.76E-08 | IFNGR1 | 4.91E-06 | ITGB3 | 5.23E-09 |
| 19 | LST1 | 1.36E-02 | TM4SF1 | 1.37E-07 | GSTA4 | 5.26E-06 | HEMGN | 6.31E-09 |
| 20 | CYBA | 1.46E-02 | NNAT | 1.37E-07 | CD86 | 5.39E-06 | PSTPIP2 | 1.01E-08 |

**Supplementary Table 7.**
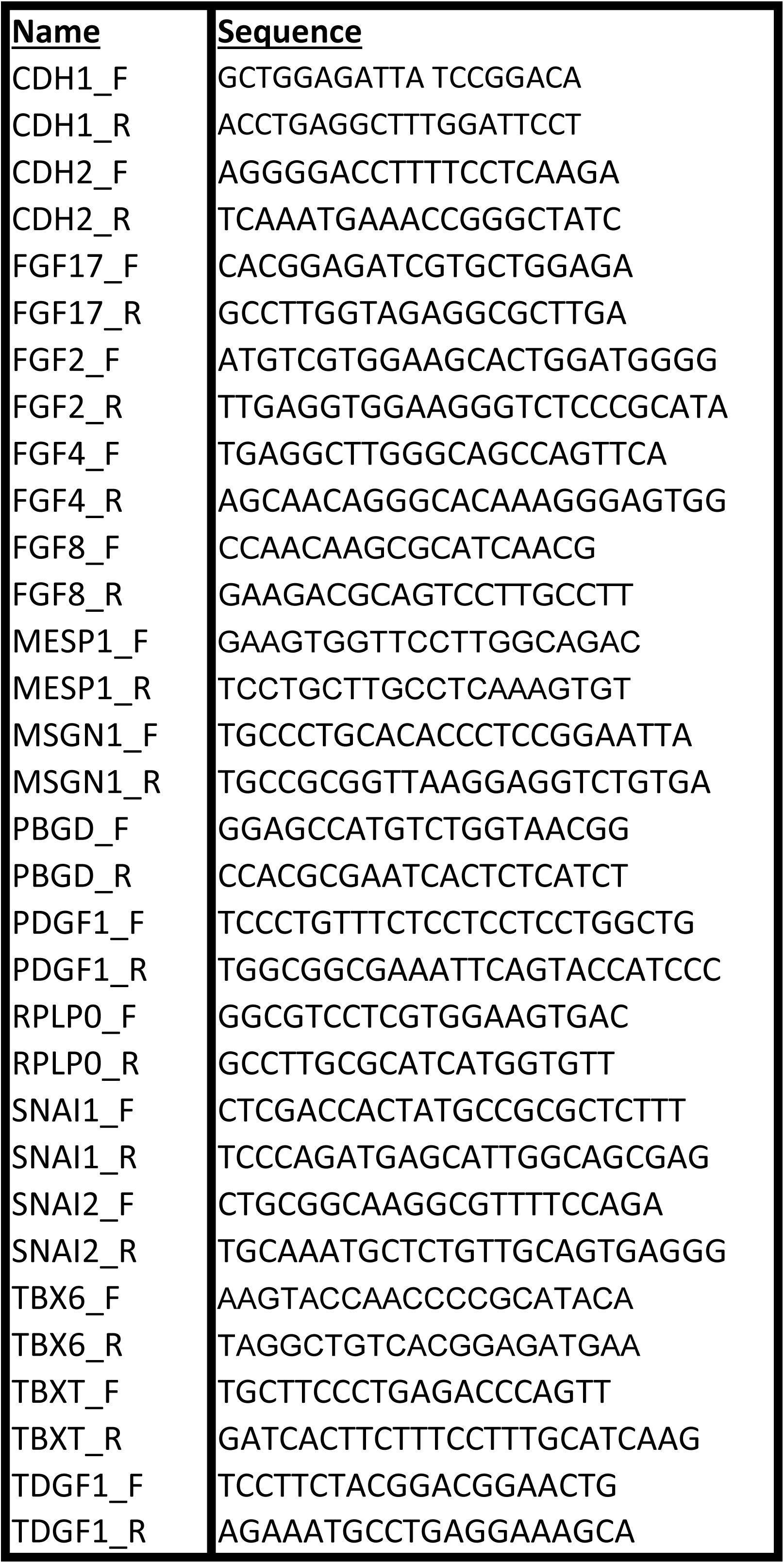
RT-PCR Primer Details.

## REFERENCES

Bergen V, Lange M, Peidli S, Wolf FA, Theis FJ. 2019. Generalizing RNA velocity to transient cell states through dynamical modeling. bioRxiv.

Bian Z, Gong Y, Huang T, Lee CZW, Bian L, Bai Z, Shi H, Zeng Y, Liu C, He J, et al. 2020. Deciphering human macrophage development at single-cell resolution. Nature.

Bianchi DW, Wilkins-Haug LE, Enders AC, Hay ED. 1993. Origin of extraembryonic mesoderm in experimental animals: Relevance to chorionic mosaicism in humans. Am J Med Genet.

Briggs JA, Weinreb C, Wagner DE, Megason S, Peshkin L, Kirschner MW, Klein AM. 2018. The dynamics of gene expression in vertebrate embryogenesis at single-cell resolution. Science (80-).

Burdsal CA, Flannery ML, Pedersen RA. 1998. FGF-2 alters the fate of mouse epiblast from ectoderm to mesoderm in vitro. Dev Biol.

Cano A, Pérez-Moreno MA, Rodrigo I, Locascio A, Blanco MJ, Del Barrio MG, Portillo F, Nieto MA. 2000. The transcription factor Snail controls epithelial-mesenchymal transitions by repressing E-cadherin expression. Nat Cell Biol.

Chalamalasetty RB, Garriock RJ, Dunty WC, Kennedy MW, Jailwala P, Si H, Yamaguchi TP. 2014. Mesogenin 1 is a master regulator of paraxial presomitic mesoderm differentiation. Dev.

Chen D, Sun N, Hou L, Kim R, Faith J, Aslanyan M, Tao Y, Zheng Y, Fu J, Liu W, et al. 2019. Human Primordial Germ Cells Are Specified from Lineage-Primed Progenitors. Cell Rep.

Chen G, Gulbranson DR, Hou Z, Bolin JM, Ruotti V, Probasco MD, Smuga-Otto K, Howden SE, Diol NR, Propson NE, et al. 2011. Chemically defined conditions for human iPSC derivation and culture. Nat Methods.

Chiquoine AD. 1954. The identification, origin, and migration of the primordial germ cells in the mouse embryo. Anat Rec.

Ciruna B, Rossant J. 2001. FGF Signaling Regulates Mesoderm Cell Fate Specification and Morphogenetic Movement at the Primitive Streak. Dev Cell.

De Bakker BS, De Jong KH, Hagoort J, De Bree K, Besselink CT, De Kanter FEC, Veldhuis T, Bais B, Schildmeijer R, Ruijter JM, et al. 2016. An interactive three-dimensional digital atlas and quantitative database of human development. Science (80-).

Delile J, Rayon T, Melchionda M, Edwards A, Briscoe J, Sagner A. 2019. Single cell transcriptomics reveals spatial and temporal dynamics of gene expression in the developing mouse spinal cord. Dev.

Ding J, Yang L, Yan YT, Chen A, Desai N, Wynshaw-Boris A, Shen MM. 1998. Cripto is required for correct orientation of the anterior-posterior axis in the mouse embryo. Nature.

Dobin A, Davis CA, Schlesinger F, Drenkow J, Zaleski C, Jha S, Batut P, Chaisson M, Gingeras TR. 2013. STAR: Ultrafast universal RNA-seq aligner. Bioinformatics.

Durinck S, Spellman PT, Birney E, Huber W. 2009. Mapping identifiers for the integration of genomic datasets with the R/ Bioconductor package biomaRt. Nat Protoc.

Efremova M, Vento-Tormo M, Teichmann SA, Vento-Tormo R. 2020. CellPhoneDB: inferring cell–cell communication from combined expression of multi-subunit ligand–receptor complexes. Nat Protoc.

Florian J, Hill JP. 1935. An Early Human Embryo (No. 1285, Manchester Collection), with Capsular Attachment of the Connecting Stalk. J Anat.

Grün D, Lyubimova A, Kester L, Wiebrands K, Basak O, Sasaki N, Clevers H, Van Oudenaarden A. 2015. Single-cell messenger RNA sequencing reveals rare intestinal cell types. Nature.

Haghverdi L, Büttner M, Wolf FA, Buettner F, Theis FJ. 2016. Diffusion pseudotime robustly reconstructs lineage branching. Nat Methods.

Irie N, Weinberger L, Tang WWC, Kobayashi T, Viukov S, Manor YS, Dietmann S, Hanna JH, Surani MA. 2015. SOX17 is a critical specifier of human primordial germ cell fate. Cell.

Jeong Kyo Yoon, Wold B. 2000. The bHLH regulator pMesogenin1 is required for maturation and segmentation of paraxial mesoderm. Genes Dev.

Jiang R, Lan Y, Norton CR, Sundberg JP, Gridley T. 1998. The slug gene is not essential for mesoderm or neural crest development in mice. Dev Biol.

Jin JZ, Ding J. 2013. Cripto is required for mesoderm and endoderm cell allocation during mouse gastrulation. Dev Biol.

Johansson BM, Wiles M V. 1995. Evidence for involvement of activin A and bone morphogenetic protein 4 in mammalian mesoderm and hematopoietic development. Mol Cell Biol.

La Manno G, Soldatov R, Zeisel A, Braun E, Hochgerner H, Petukhov V, Lidschreiber K, Kastriti ME, Lönnerberg P, Furlan A, et al. 2018. RNA velocity of single cells. Nature.

Lawson KA, Wilson V. 2016. A Revised Staging of Mouse Development Before Organogenesis. In Kaufman’s Atlas of Mouse Development Supplement.

Leng N, Chu LF, Barry C, Li Y, Choi J, Li X, Jiang P, Stewart RM, Thomson JA, Kendziorski C. 2015. Oscope identifies oscillatory genes in unsynchronized single-cell RNA-seq experiments. Nat Methods.

Li H. 2013. [Heng Li - Compares BWA to other long read aligners like CUSHAW2] Aligning sequence reads, clone sequences and assembly contigs with BWA-MEM. arXiv Prepr arXiv.

Love MI, Huber W, Anders S. 2014. Moderated estimation of fold change and dispersion for RNA-seq data with DESeq2. Genome Biol.

Lun ATL, Bach K, Marioni JC. 2016. Pooling across cells to normalize single-cell RNA sequencing data with many zero counts. Genome Biol.

Magnúsdóttir E, Azim Surani M. 2014. How to make a primordial germ cell. Dev.

Martyn I, Kanno TY, Ruzo A, Siggia ED, Brivanlou AH. 2018. Self-organization of a human organizer by combined Wnt and Nodal signaling. Nature.

McGrath KE, Frame JM, Palis J. 2015. Early hematopoiesis and macrophage development. Semin Immunol.

Mendjan S, Mascetti VL, Ortmann D, Ortiz M, Karjosukarso DW, Ng Y, Moreau T, Pedersen RA. 2014. NANOG and CDX2 pattern distinct subtypes of human mesoderm during exit from pluripotency. Cell Stem Cell.

Meno C, Gritsman K, Ohishi S, Ohfuji Y, Heckscher E, Mochida K, Shimono A, Kondoh H, Talbot WS, Robertson EJ, et al. 1999. Mouse lefty2 and zebrafish antivin are feedback inhibitors of nodal signaling during vertebrate gastrulation. Mol Cell.

Messmer T, von Meyenn F, Savino A, Santos F, Mohammed H, Lun ATL, Marioni JC, Reik W. 2019. Transcriptional Heterogeneity in Naive and Primed Human Pluripotent Stem Cells at Single-Cell Resolution. Cell Rep.

Moris N, Anlas K, van den Brink SC, Alemany A, Schröder J, Ghimire S, Balayo T, van Oudenaarden A, Martinez Arias A. 2020. An in vitro model of early anteroposterior organization during human development. Nature.

Nieto MA, Sargent MG, Wilkinson DG, Cooke J. 1994. Control of cell behavior during vertebrate development by Slug, a zinc finger gene. Science (80-).

Nowotschin S, Ferrer-Vaquer A, Concepcion D, Papaioannou VE, Hadjantonakis AK. 2012. Interaction of Wnt3a, Msgn1 and Tbx6 in neural versus paraxial mesoderm lineage commitment and paraxial mesoderm differentiation in the mouse embryo. Dev Biol.

Nowotschin S, Setty M, Kuo YY, Liu V, Garg V, Sharma R, Simon CS, Saiz N, Gardner R, Boutet SC, et al. 2019. The emergent landscape of the mouse gut endoderm at single-cell resolution. Nature.

O’Rahilly R, Müller F. 2010. Developmental stages in human embryos: Revised and new measurements. Cells Tissues Organs.

O’Rahilly R, Müller F. 1987. Developmental Stages in Human Embryos. Contrib Embryol, Carnegie Inst Wash 637.

Ornitz DM, Itoh N. 2015. The fibroblast growth factor signaling pathway. Wiley Interdiscip Rev Dev Biol.

Ortega S, Ittmann M, Tsang SH, Ehrlich M, Basilico C. 1998. Neuronal defects and delayed wound healing in mice lacking fibroblast growth factor 2. Proc Natl Acad Sci U S A.

Palis J. 2016. Hematopoietic stem cell-independent hematopoiesis: emergence of erythroid, megakaryocyte, and myeloid potential in the mammalian embryo. FEBS Lett.

Patro R, Duggal G, Love MI, Irizarry RA, Kingsford C. 2017. Salmon provides fast and bias-aware quantification of transcript expression. Nat Methods.

Peng G, Suo S, Chen J, Chen W, Liu C, Yu F, Wang R, Chen S, Sun N, Cui G, et al. 2016. Spatial Transcriptome for the Molecular Annotation of Lineage Fates and Cell Identity in Mid-gastrula Mouse Embryo. Dev Cell.

Petropoulos S, Edsgärd D, Reinius B, Deng Q, Panula SP, Codeluppi S, Plaza Reyes A, Linnarsson S, Sandberg R, Lanner F. 2016. Single-Cell RNA-Seq Reveals Lineage and X Chromosome Dynamics in Human Preimplantation Embryos. Cell.

Picelli S, Faridani OR, Björklund ÅK, Winberg G, Sagasser S, Sandberg R. 2014. Full-length RNA-seq from single cells using Smart-seq2. Nat Protoc.

Pijuan-Sala B, Griffiths JA, Guibentif C, Hiscock TW, Jawaid W, Calero-Nieto FJ, Mulas C, Ibarra-Soria X, Tyser RCV, Ho DLL, et al. 2019. A single-cell molecular map of mouse gastrulation and early organogenesis. Nature 566: 490–495.

Sasaki K, Nakamura T, Okamoto I, Yabuta Y, Iwatani C, Tsuchiya H, Seita Y, Nakamura S, Shiraki N, Takakuwa T, et al. 2016. The Germ Cell Fate of Cynomolgus Monkeys Is Specified in the Nascent Amnion. Dev Cell.

Scialdone A, Natarajan KN, Saraiva LR, Proserpio V, Teichmann SA, Stegle O, Marioni JC, Buettner F. 2015. Computational assignment of cell-cycle stage from single-cell transcriptome data. Methods.

Segal JM, Kent D, Wesche DJ, Ng SS, Serra M, Oulès B, Kar G, Emerton G, Blackford SJI, Darmanis S, et al. 2019. Single cell analysis of human foetal liver captures the transcriptional profile of hepatobiliary hybrid progenitors. Nat Commun.

Simunovic M, Metzger JJ, Etoc F, Yoney A, Ruzo A, Martyn I, Croft G, You DS, Brivanlou AH, Siggia ED. 2019. A 3D model of a human epiblast reveals BMP4-driven symmetry breaking. Nat Cell Biol.

Smith DE, Franco Del Amo F, Gridley T. 1992. Isolation of Sna, a mouse gene homologous to the Drosophila genes snail and escargot: Its expression pattern suggests multiple roles during postimplantation development. Development.

Stern CD. 2004. Gastrulation: From Cells to Embryo.

Streit A. 2007. The preplacodal region: An ectodermal domain with multipotential progenitors that contribute to sense organs and cranial sensory ganglia. Int J Dev Biol.

Stuart T, Butler A, Hoffman P, Hafemeister C, Papalexi E, Mauck WM, Hao Y, Stoeckius M, Smibert P, Satija R. 2019. Comprehensive Integration of Single-Cell Data. Cell.

Sun X, Meyers EN, Lewandoski M, Martin GR. 1999. Targeted disruption of Fgf8 causes failure of cell migration in the gastrulating mouse embryo. Genes Dev.

Teo AKK, Arnold SJ, Trotter MWB, Brown S, Ang LT, Chng Z, Robertson EJ, Dunn NR, Vallier L. 2011. Pluripotency factors regulate definitive endoderm specification through eomesodermin. Genes Dev.

Thisse B, Thisse C. 2005. Functions and regulations of fibroblast growth factor signaling during embryonic development. Dev Biol.

Trevers KE, Prajapati RS, Hintze M, Stower MJ, Strobl AC, Tambalo M, Ranganathan R, Moncaut N, Khan MAF, Stern CD, et al. 2017. Neural induction by the node and placode induction by head mesoderm share an initial state resembling neural plate border and ES cells. Proc Natl Acad Sci U S A.

Wagner DE, Weinreb C, Collins ZM, Briggs JA, Megason SG, Klein AM. 2018. Single-cell mapping of gene expression landscapes and lineage in the zebrafish embryo. Science (80-).

Warmflash A, Sorre B, Etoc F, Siggia ED, Brivanlou AH. 2014. A method to recapitulate early embryonic spatial patterning in human embryonic stem cells. Nat Methods.

Yamaguchi Y, Yamada S. 2019. The kyoto collection of human embryos and fetuses: History and recent advancements in modern methods. Cells Tissues Organs.

Yang L, Zhang H, Hu G, Wang H, Abate-Shen G, Shen MM. 1998. An early phase of embryonic Dlx5 expression defines the rostral boundary of the neural plate. J Neurosci.

Yang R, Van Etten JL, Dehm SM. 2018. Indel detection from DNA and RNA sequencing data with transIndel. BMC Genomics.

Yiangou L, Grandy RA, Morell CM, Tomaz RA, Osnato A, Kadiwala J, Muraro D, Garcia-Bernardo J, Nakanoh S, Bernard WG, et al. 2019. Method to Synchronize Cell Cycle of Human Pluripotent Stem Cells without Affecting Their Fundamental Characteristics. Stem Cell Reports.

Zhou M, Sutliff RL, Paul RJ, Lorenz JN, Hoying JB, Haudenschild CC, Yin M, Coffin JD, Kong L, Kranias EG, et al. 1998. Fibroblast growth factor 2 control of vascular tone. Nat Med.

